# Loss of Insulin Signaling in Microglia Impairs Cellular Uptake of Aβ and Neuroinflammatory Response Exacerbating Alzheimer-like Neuropathology

**DOI:** 10.1101/2024.08.22.609112

**Authors:** Wenqiang Chen, Xiangyu Liu, Vitor Rosetto Munoz, C. Ronald Kahn

## Abstract

Insulin receptors are present on cells throughout the body, including the brain. Dysregulation of insulin signaling in neurons and astrocytes has been implicated in altered mood, cognition, and the pathogenesis of Alzheimer’s disease (AD). To define the role of insulin signaling in microglia, the primary phagocytes in brain critical for maintenance and damage repair, we created mice with an inducible microglia-specific insulin receptor knockout (MG-IRKO). RiboTag profiling of microglial mRNAs revealed that loss of insulin signaling results in alterations of gene expression in pathways related to innate immunity and cellular metabolism. *In vitro*, loss of insulin signaling in microglia results in metabolic reprograming with an increase in glycolysis and impaired uptake of Aβ. *In vivo*, MG-IRKO mice exhibit alterations in mood and social behavior, and when crossed with the 5xFAD mouse model of AD, the resultant mice exhibit increased levels of Aβ plaque and elevated neuroinflammation. Thus, insulin signaling in microglia plays a key role in microglial cellular metabolism, neuroinflammation and the ability of the cells to take up Aβ, such that reduced insulin signaling in microglia alters mood and social behavior and accelerates AD pathogenesis. Together these data indicate key roles of insulin action in microglia and the potential of targeting insulin signaling in microglia in treatment of AD.

## Introduction

Type 2 diabetes (T2D), obesity and metabolic syndrome are major causes of morbidity and mortality worldwide. In all of these disorders, insulin resistance is a major pathophysiological feature. Growing evidence has shown that insulin resistance in these disorders is also associated with increased risks of brain disorders, including Alzheimer’s disease (AD) and depression (Barbagallo and Dominguez, 2014; Chen et al., 2022; Talbot et al., 2012). The cellular and molecular factors that contribute to these neuropsychiatric disorders are complex, and exactly how dysregulated insulin signaling might contribute to this comorbidity remains incompletely understood (Frolich et al., 1998; Talbot *et al*., 2012). Studies have suggested that insulin resistance can alter brain function and development of AD through multiple mechanisms. These include alterations in brain function at the cellular level, as well as via vascular dysfunction and inflammation (Arnold et al., 2018; Kellar and Craft, 2020). Previously, we have used mice with insulin receptor (IR) knockout in whole brain (NIRKO mice) to explore this interaction and demonstrated that these mice not only exhibit mild metabolic syndrome (Bruning et al., 2000), but also show mood and behavioral alterations and increased tau phosphorylation, a hallmark of AD (Schubert et al., 2004). More recently we found that loss of insulin signaling in astrocytes can also contribute to mood disturbances by affecting astrocyte ATP release which secondarily reduces neuronal dopamine release (Cai et al., 2018), and that when crossed with an AD mouse model, mice lacking insulin signaling in astrocytes exhibited exacerbation of AD phenotypes (Chen et al., 2023).

Microglia serve as the brain-resident immune cells and have been shown to play important roles in AD pathogenesis through their many functions, including cytokine release, phagocytosis, and synapse pruning (Keren-Shaul et al., 2017; Paolicelli et al., 2022). Intranasal insulin administration has also shown some beneficial effects in human and rodent models of AD, partially through increasing Aβ clearance and alleviating aberrant microglia activation (Chen et al., 2014; Frazier et al., 2020; Wong et al., 2024). Whether insulin resistance in microglia plays a role in brain function and development of AD remains unknown.

To understand the role of insulin signaling in microglia and how dysregulated insulin action in microglia might contribute to the comorbidity of T2D and AD, in the present study, we have created mice with microglia insulin resistance by knockout of the IR using a mouse carrying a TMEM119^CreERT2^ transgene that specifically targets brain parenchymal microglia, but not other brain macrophages (Kaiser and Feng, 2019). We found that mice with microglia-specific IR knockout develop minimal alterations in systemic metabolism, but significant changes in mood and social behaviors. At the cellular level, we find that insulin signaling regulates microglial cellular metabolism, the ability of microglia to take up Aβ and the development of neuroinflammatory responses in the brain. As a result, loss of IR signaling in microglia exacerbates Aβ deposition in brain areas associated with cognition. Thus, insulin signaling in microglia plays key roles in regulation of neuroinflammation and Aβ uptake, driving the comorbidity of T2D and AD and providing a novel target for modulating disease development.

## Results

### Deletion of InsR in Microglia

To study the role of insulin receptor signaling in microglia, we generated mice with selective deletion of *Insr* in microglia (MG-IRKO, Figure 1A) by crossing mice carrying conditional alleles of *Insr* (*Insr*-floxed mice, IR^f/f^) (Bruning et al., 1998) with mice expressing the tamoxifen-inducible Cre recombinase (Cre^ERT2^) transgene driven by the Tmem119 promoter (TMEM119^CreERT2^) (Bennett et al., 2016; Kaiser and Feng, 2019). Previous studies have shown that TMEM119 is specifically expressed in microglia in both humans and rodents, but not in other brain-resident cells nor infiltrating macrophages (Bennett *et al*., 2016; Ruan et al., 2020; Satoh et al., 2016). To avoid any developmental effects, we administrated tamoxifen to all mice starting at age of 6 weeks. Behavioral and metabolic characterization was performed at least 4 weeks after the induction of IR knockout in microglia.

**Figure 1.**
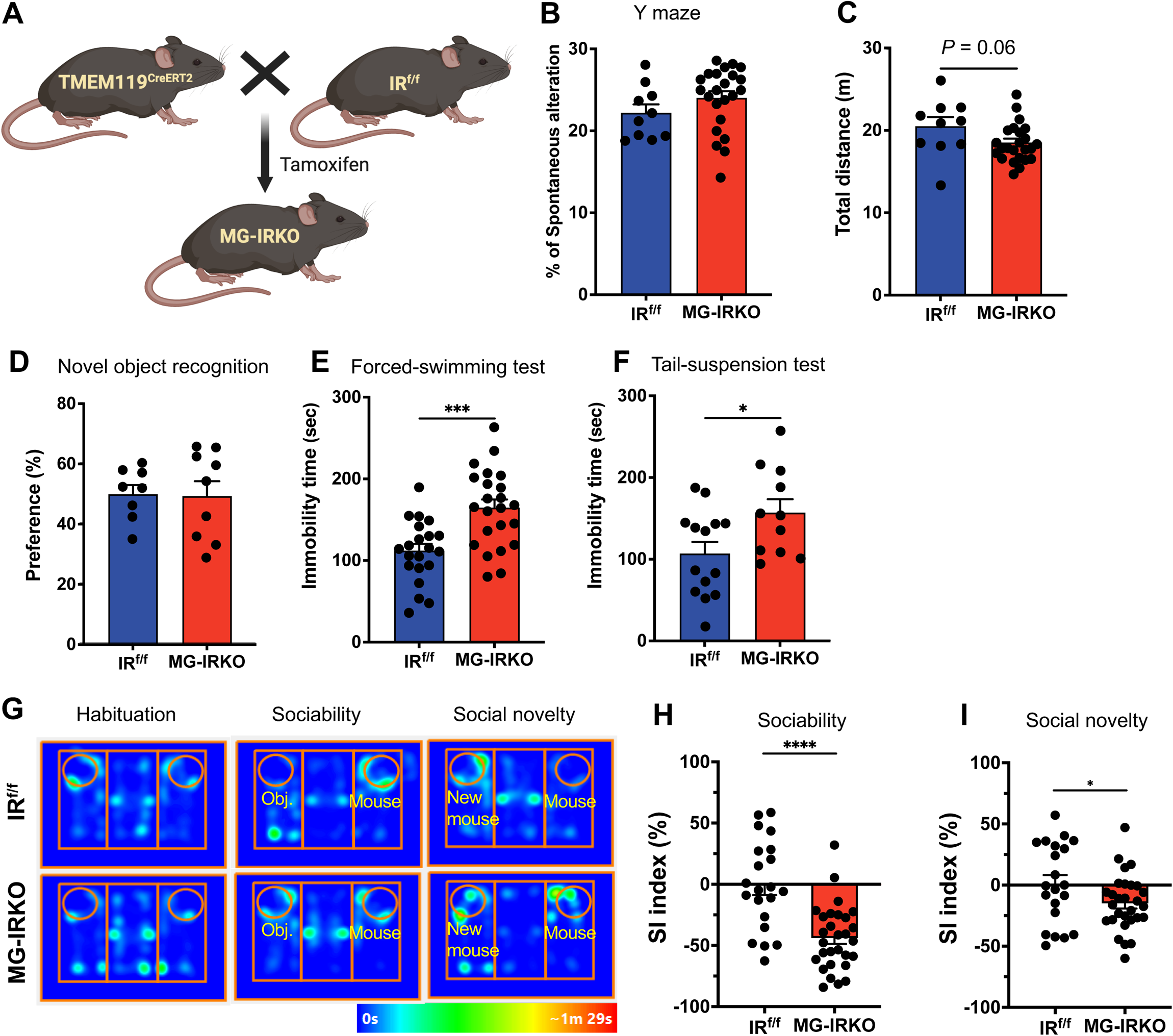
Effects of Microglia IRKO on behavioral performances. (A) Schematic showing the stragety for generating mice with selective deletion of *Insr* in microglia. (B-C) Y maze performance of 3-month IR^f/f^ and MG-IRKO male mice, showing (B) percentage of spontaneous alteration (%) and (C) total distance (m). N = 10-24; (D) Percentage of preference score during the novel object recognition of 3-month IR^f/f^ and MG-IRKO male mice N = 8-9; (E) Immobility time of 3-month IR^f/f^ and MG-IRKO male mice in the forced-swimming test. N = 21-24; (F) Immobility time of 3-month IR^f/f^ and MG-IRKO male mice in the tail-suspension test. N = 11-14; (G) Representative images showing heatmap of duration in each chamber of 3-month IR^f/f^ and MG-IRKO male mice in the three-chamber social interaction test. N = 22-29. Obj., object. (H-I) Sociability index (SI) of 3-month IR^f/f^ and MG-IRKO male mice in (H) sociability session and (I) social novelty session in three -chamber social interaction test. Data are mean ± SEM. \**P* < 0.05, \*\*\**P* < 0.001, \*\*\*\**P* < 0.0001, unpaired *t*-test.

To confirm the efficiency of IR knockout, we isolated CD11b^+^ microglia from dispersed whole brain cell mixtures using Dynabeads and biotinylated anti-CD11b antibody, as previously described (Kim et al., 2019). RT-qPCR of the isolated in CD11b^+^ cells revealed a 10- to 120-fold enrichment of microglia signature genes, including *Cd83*, *Hexb*, *Trem2*, *Fcrls*, *Hpgd*, *P2ry12*, and *Grp34* as compared to the CD11b^-^ cells (Figure S1A). Conversely, expression of marker genes for other glial cell types, such as astrocytes (*Aldh1l1*, *Aldoc*, and *S100b*) and oligodendrocytes (*Mbp* and *Sox10*), were depleted in CD11b^+^ cells (Figure S1B), demonstrating the validity of the microglial isolation protocol. RT-qPCR in isolated CD11b^+^ cells of MG-IRKO brains revealed a >75% decrease in *Insr* mRNA as compared to CD11b^+^ cells of the IR^f/f^ brains, while analysis of the unfractionated cortex revealed no difference in expression of *Insr* gene, indicating the specificity of the knockout (Figure S1C). Thus, the TMEM119 promoter-driven *Insr* deletion was specific to microglia, and when coupled with a tamoxifen-inducible Cre, can achieve a robust reduction of *Insr* expression in microglial cells in the brain.

### Deletion of IR in microglia produces minimal alteration in systemic metabolism

To determine whether loss of IR signaling in microglia produces any metabolic phenotypes, we subjected the MG-IRKO mice to a series of metabolic examinations. DEXA scan revealed no significant differences in fat mass, lean mass, or distribution of fat in MG-IRKO mice versus controls (Figure S2A). There was no change in serum insulin in male MR-IRKO mice, but there was a small, but significant, increase in fasting serum insulin levels in female MR-IRKO mice (Figure S2B). Serum IGF1 levels were unchanged in both sexes (Figure S2C). IP glucose tolerance tests in both male and female MG-IRKO mice revealed no major change in glucose tolerance (Figures S2D-2G), but both sexes showed slightly improved insulin sensitivity during an IP-insulin tolerance test, and this reached significance in male mice (Figure S2F). CLAMS metabolic cage assessment of 4-month-old mice revealed no significant differences in food or water intake (Figures S3A-3D), O_2_ consumption, or CO_2_ production in either male or female MG-IRKO mice (Figures S3E and 3F). Although TMEM119 is known to be expressed in bone, and previous studies have suggested a role in osteoblast differentiation and bone development (Jiang et al., 2017; Tanaka et al., 2012), MG-IRKO mice showed no differences in bone mineral density or bone mineral content as measured by DEXA scan (Figures S3G and 3H). Thus, deletion of IR in microglia produces minimal alteration in systemic metabolism.

### Effects of Microglia IRKO on behavioral performances

To determine whether microglia-specific IR deletion leads to behavioral alterations, we performed a battery of tests assessing anxiety and cognition in mice of both sexes. MG-IRKO mice showed no differences in anxiety-like phenotype, as measured by the light-dark transition test (Figures S4A and 4B), no difference in total sucrose intake or sucrose preference (Figure S4C), and no difference in working memory as assessed by the Y-maze, although MG-IRKO mice showed a trend to a decrease in total distance traveled (Figure 1B and 1C). Likewise, the novel object recognition test revealed no preferences in time spent on novel object (Figure 1D and Figure S4D), and both groups of mice showed similar performance in the marble burying test (Figure S4E). By contrast, in both the forced-swimming test (FST) and the tail-suspension test (TST), two widely used assays to assess depressive or stress-coping behaviors in rodents (Castagne et al., 2011), MG-IRKO mice showed ∼1.5-fold increases in immobility time compared to the controls (*P* < 0.001 and *P* < 0.05, Figure 1E and 1F, respectively). Thus, loss of microglia IR signaling leads to depressive-like phenotypes, but no change in anxiety, locomotion, or working memory.

To determine if loss of IR signaling in microglia modifies sociability, we performed a three-chamber social interaction (SI) test across three sessions: habituation, familiarization, and social novelty (Figure 1G). During the habituation session, MG-IRKO mice had a significantly decreased locomotion (Figure S4F) and a trend of increase in immobility time (*P* = 0.06, Figure S4G). More importantly, in the social novelty sessions, these mice exhibited a significant increase in immobility time (Figure S4G), a phenotype of social defeat. In addition, whereas MG-IRKO mice spent equal amounts of time in the two empty chambers during the habituation session (Figure S4H), when exposed to a stranger mouse during the familiarization session, MG-IRKO mice spent significantly less time in the chamber with the stranger mouse (*P* < 0.0001, Figure S4I), indicating social avoidance. Likewise, MG-IRKO mice had significantly decreased social interaction during the familiarization session (*P* < 0.0001, Figure 1H). In the social novelty session, when an inanimate object was replaced with a novel mouse, MG-IRKO mice spent more time with the mouse that had been interacted during the previous session, less time in the neutral middle chamber (Figure S4J) and a similar amount of time with the novel mouse. As a result, the social interaction (SI) score was significantly lower in the MG-IRKO mice than in controls (*P* < 0.05, Figure 1I). Taken together, these data demonstrate that MG-IRKO mice exhibit impaired social novelty preference, i.e., an autism-like behavior, as well as depressive-like phenotypes.

### RiboTag profiling reveals pathways affected by microglia IR deletion

To begin to determine the molecular mechanisms by which IR deletion in microglia can affect behavior, we crossed MG-IRKO mice with RiboTag mice (homozygous for the Rpl22^HA^ allele) to create MG-IRKO^RiboTag^ mice (Sanz et al., 2019; Sanz et al., 2009). This enables immunoprecipitation (IP) of microglial-specific ribosome-bounded mRNA from whole brain extracts and assessment of the effect of loss of insulin action on gene expression in microglia (Figure 2A). RiboTag mice with Cre recombinase expression without the flox alleles were used as controls (MG^RiboTag^). In control RiboTag mice, RT-qPCR analysis for microglia marker gene *Tmem119* revels an over 30-fold enrichment of *Tmem119* in the RiboTag *IP* and a parallel 55% depletion in the supernatant fraction (Figure 2B), demonstrating an efficient tagging of microglia-bounded ribosomes.

**Figure 2.**
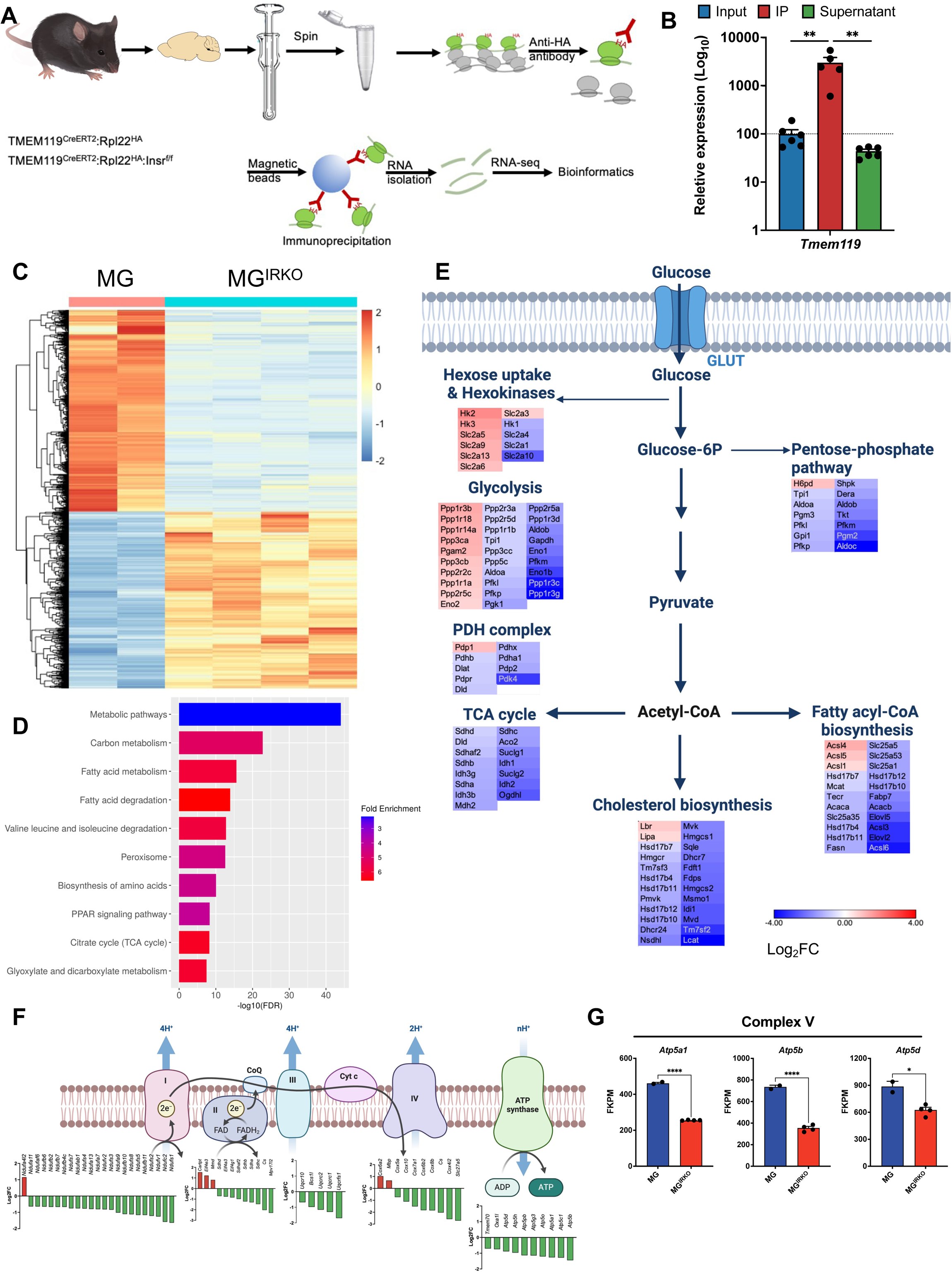
RiboTag profiling reveals pathways affected by microglia IR deletion. (A) Schematic showing procedure of RiboTag profiling. (B) RT-qPCR analysis of mRNA extracted from inputs, IP, and supernatant fraction in brains of MG-RiboTag mouse (N = 4-6). Data are mean ± SEM. \*\**P* < 0.01, unpaired *t*-test, on log_10_ scale. (C) Heatmap showing down- and up-regulated DEGs, FDR = 0.01, fold change (FC) = 2. (D) KEGG pathway analysis showing down-regulated DEGs were related to cellular metabolism. (E) Heatmap showing regulation of genes involved in glucose metabolism, including hexose uptake and hexokinase, glycolysis, pentose phosphate pathway, PDH complex, TCA cycle, fatty acyl-CoA biosynthesis, and cholesterol biosynthesis. The magitude of regulation is represented in the heatmap by log_2_FC value (MR-IRKO/ IR^f/f^ control). (F) Upper: Schematic showing mitochondrial respiratory chain complex I – V. Lower: DEGs involved in the regulation of each complex are represented by Log_2_FC. (G) FKPM values of example list of genes encoding pathways of mitochondrial complex V. N = 2-4. Data are mean ± SEM. \**P* < 0.05, \*\*\*\**P* < 0.0001, unpaired *t*-test.

As expected, comparison of RiboTagged mRNAs from *IP* samples versus *Input* samples from the control mice revealed a strong enrichment of multiple microglial genes including *Tmem119*, *Hexb*, *Tyrobp*, validating the efficiency of RiboTag profiling. Comparing *IP* samples from MG-IRKO^RiboTag^ and MG^RiboTag^ mice revealed 6,394 differentially expressed genes (DEG) with a fold change (FC) ≥ │1.5│ and *P* < 0.01 (Figure 2C), reflecting the transcriptional alterations in microglia following IR deletion. Of these, 2,658 genes were upregulated, and 3,736 genes were downregulated (Figure 2C). KEGG enrichment analysis of the significantly decreased DEGs revealed that many of these were in pathways associated with cellular metabolism, including carbon metabolism, fatty acid metabolism, branched chain amino acid metabolism, as well as TCA cycle (Figure 2D), whereas the significantly upregulated genes were in pathways associated with phagocytosis, chemokine and cytokine pathways (Figure 3A).

**Figure 3.**
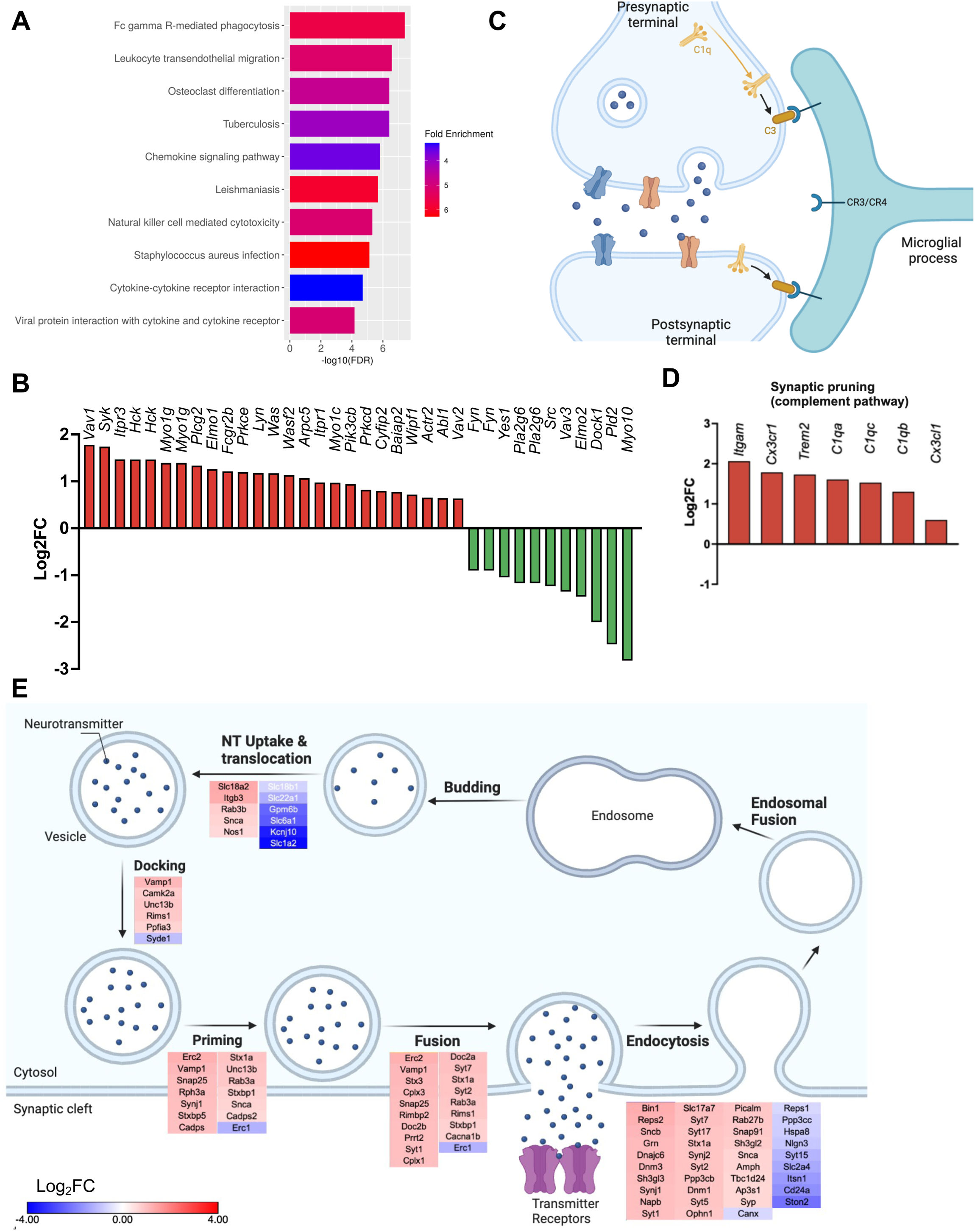
Microglia IR deletion impairs microglia pathways. (A) KEGG pathway analysis showing up-regulated DEGs related to innate immune pathways, including phagocytosis, chemokine and cytokine signaling. (B) DEGs involved in the regulation of Fc receptor-mediated phagocytosis are represented by Log_2_FC. (C) Schematic showing microglia complement process. (D) DEGs involved in the regulation of synaptic pruning. (E) Schematic and heatmap showing DEGs involved in the regulation of synaptic vesicle recycling. The magitude of regulation is represented in the heatmap by log_2_FC value (MR-IRKO / IR^f/f^ control).

Using brain samples (including the hypothalamus, hippocampus and nucleus accumbens) from mice undergoing a euglycemic, hyperinsulinemic clamp, we previously demonstrated that insulin can regulate genes of glucose uptake and metabolism (Cai et al., 2021). Consistent with this, we found that loss of IR signaling in microglia caused a downregulation of multiple genes in various steps in intracellular glucose metabolism, including hexose uptake and hexokinases, glycolysis, PDH complex (Figure 2E). However, interestingly, some of genes in hexose phosphorylation (*Hk2* and *Hk3*) and glycolysis (*Ppp1r3b* and *Eno2*) showed upregulation (Figure 2E). In addition, IRKO in microglia caused a broad downregulation of genes in mitochondrial complexes I-V and genes of oxidative phosphorylation (Figure 2F). Some of the most striking examples were genes encoding mitochondrial complex II (*Shda*, *Shdb*, *Shdc*) and complex V (*Atp5a1*, *Atp5b*, *Atp5d*), which exhibited a 50% or more reduction (Figure 2G and Figure S5A). On the other hand, KEGG analysis of the significantly upregulated genes following microglial IR deletion revealed that most were associated with immune activation, including Fc receptor-mediated phagocytosis, chemokine signaling pathway, and cytokine-cytokine receptor interaction (Figure 3A-B). Specific examples of these include phagocytosis-promoting genes (*Eif5a*, *Uchl1*, *Tubb5*, Figure S6A), chemokine and cytokine genes (*Cxcl12*, *Cxcl16*, *Ifngr1*, *Ifngr2*, *Cx3cr1*, Figure S6B), a number of which increased by 3-fold or more.

Microglia are known to play an essential role in the regulation of synaptic pruning and synaptic plasticity (Mordelt and de Witte, 2023). Dysfunction of these processes during neurodevelopment have been associated with schizophrenia and autism spectrum disorders (Sellgren et al., 2019; Wu et al., 2024). Given that microglia IR deletion in mice produces an autism-like phenotype as revealed by the social interaction test (Figure 1G and 1I), we specifically analyzed the gene pathways related to synaptic pruning and synaptic vesicle recycling. Our analysis shows that IR deletion in microglia enhanced expression of multiple complement genes, including *C1qa*, *C1qb*, and *C1qc*, by 2- to 4-folds (Figures 3C and 3D). Also, consistent with the alterations in the behavioral tests, we found that key genes encoding proteins involved in presynaptic vesicle recycling, neurotransmitter uptake, vesicle fusion, and endocytosis were significantly upregulated after microglia IRKO (Figure 3E). These included a nearly 3-fold increase in gene expression of *Slc17a6*, *Slc18a2*, *Bin1*, *Erc2*, and *Vamp1*. Taken together, our data reveal that microglia IR deletion produces profound effects on expression of genes involved in cellular metabolism and microglia immune activation, with former being mainly down-regulated and the latter mainly up-regulated.

### Microglia IR KO increases mitochondrial fission and impairs mitochondrial metabolism

Insulin inhibits autophagy in classic target tissues such as liver, muscle and fat (Frendo-Cumbo et al., 2021), as well as astrocytes (Geffken et al., 2022), and thus loss of insulin signaling in in these tissues increases autophagy and also the closely associated mitophagy process (Chen *et al*., 2023; Frendo-Cumbo *et al*., 2021). RiboTag profiling revealed that expression of mitophagy genes *Bnip3*, *Lrrk1*, *Bax*, *Hk2*, and *Fis1* was increased by 1.5- to 5-fold in MG-IRKO mice, whereas expression of *Pink1*, a negative regulator of autophagy and mitophagy, showed 2.7-fold decrease (Figure S7A), indicating that loss of IR signaling in microglia increases the level of mitophagy. Many of the DEGs in the autophagy and mitophagy pathways are also involved in the processes of formation of phagophore, autolysosome organization and SNARE complex (Figure S7B). Transmission Electron Microscopy (TEM) analysis of mitochondrial morphology in microglia in the hypothalamus revealed that IR deletion resulted in significantly decreased mitochondrial size, both in area and perimeter (Figure 4A-D), with no change in shape (roundness/circularity, Figure S8A and S8B). These data are consistent with the changes in gene expression revealed by the RiboTag profiling (Figure S5B) and suggest increased mitochondrial fission/mitophagy following IR deletion in microglia.

**Figure 4.**
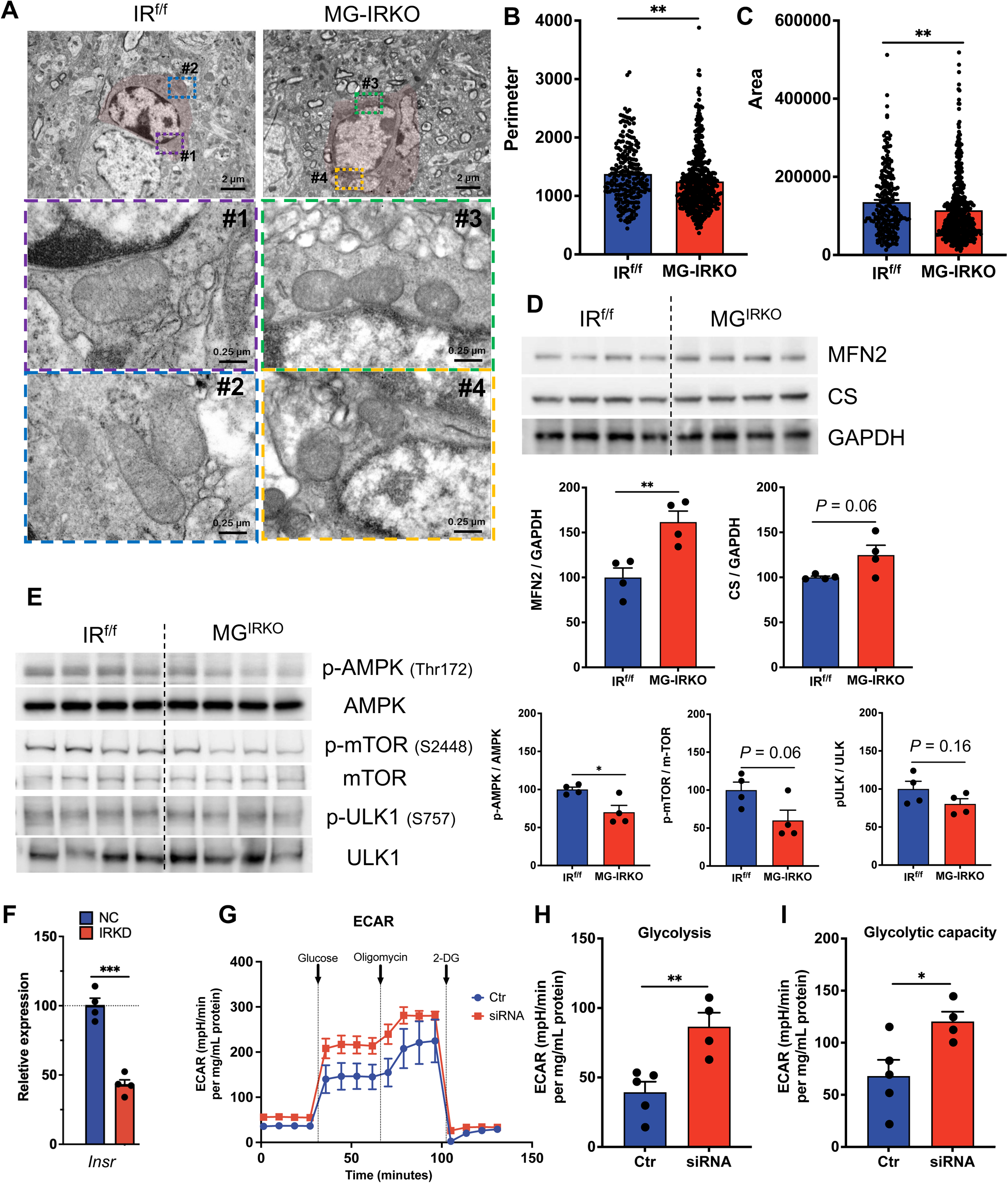
Loss of microglia IR signaling impairs cellular metabolism. (A) Representative TEM images showing mitochondrial morphology in microglia located in the ventral hypothalamus from IR^f/f^ and MG-IRKO mice (4-month-old, male). Respective enlarged images were numbered. (B-C) TEM analysis of mitochondria morphology. Bar graph showing mitochondrial area (B) and perimeter (C). N = 48 microglia from N = 4 IR^f/f^ mice, and N = 57 microglia from N = 4 MG-IRKO mice. (D) Immunoblot analysis of MFN2 and CS proteins in brain hippocampal homoganates from IR^f/f^ and MG-IRKO mice (4-month-old, male). N = 4. (E) Immunoblot analysis of phosphorated AMPK, mTOR, and ULK1 proteins and their total proteins in brain hippocampal homoganates from IR^f/f^ and MG-IRKO mice (4-month-old, male). The protein levels were normalized to GAPDH or respective total proteins. N = 4. (F) qPCR analysis of relative mRNA expression of *Insr* in microglial cells with IR KD. N = 4. unpaired *t*-test. (G-I) Seahorse analysis of IRKD microglia subjected to glycolysis stress, showing ECAR curve (G), glycolysis (H) and glycolytic capacity (I). N = 4-5. Data are mean ± SEM. \**P* < 0.05, \*\**P* < 0.01, \*\*\**P* < 0.001, unpaired *t*-test (B-F) or two-way RM ANOVA (G-I).

To better understand the consequences of microglia IRKO on mitochondrial regulation in the brain, we performed immunoblot analyses on homogenates from the hippocampus of 4-month-old control and MG-IRKO mice for markers of mitochondria and mitophagy. Expression of markers for mitochondrial complexes at the whole-tissue level was not changed in both IR^f/f^ and MG-IRKO groups (Figure S8C), and some autophagy markers remained similar levels including LC3-I, p62 and ATG (Figure S8D). There was also a 60% increase in mitofusin 2 (MFN2, Figure 4D) and a trend of increase in citrate synthase (CS), a mitochondrial enzyme indicative of mitochondrial integrity (Chhimpa et al., 2023). Phosphorylation of mitochondrial fission protein DRP1 (Dynamin-related protein 1) showed a 25% decrease at Ser^616^, but not at Ser^637^ (Figure S8E). There was no difference in whole-tissue level of expression of the mitophagy regulators Fis1 and Parkin (Figure S8F).

The AMP-activated protein kinases (AMPK) can phosphorylate proteins important for cellular metabolism, autophagy and mitochondrial functions (Steinberg and Carling, 2019). The AMPK protein interacts with the ULK1 protein kinase to initiate autophagy (Park et al., 2023; Roach, 2011), however, in insulin resistant states, AMPK activity can be diminished (Xu et al., 2012). Western blot analysis of the hippocampal homogenates revealed that phosphorylation of AMPKα at Thr^172^, a major site of activation, was significantly decreased in MG-IRKO brains (Figure 4E), whereas phosphorylation of ULK1 at Ser^757^, a site phosphorylation involved in inactivation, showed a trend of decrease (Figure 4E). The mammalian target of rapamycin (mTOR) can act as an autophagy inhibitor by phosphorylating ULK1 at Ser^757^ and disrupting the ULK1/AMPK interaction (Alers et al., 2012; Kim et al., 2011). In MG-IRKO mice, phosphorylation of mTOR at Ser^2448^ was decreased (*P* = 0.06, Figure 4E), suggesting an mTOR-activity and mTOR-dependent ULK1 inactivation. Taken together, these findings demonstrate that loss of insulin signaling in microglia increases mitochondrial fission and alters the autophagy/mitophagy pathways in these cells.

To determine the functional consequences of these changes in mitochondria, we performed knockdown of the *Insr* using siRNA in SIM-A9 cells, an immortalized murine microglia cell line (Nagamoto-Combs et al., 2014). Treatment with siRNA resulted in a 50–60% decrease of *Insr* mRNA (*P* < 0.001, Figure 4F). Previous studies have demonstrated that microglia can switch from mitochondrial oxidative phosphorylation in the unstimulated state to a glycolytic metabolism following microglial activation (Orihuela et al., 2016; Wang et al., 2019). To determine whether loss of insulin signaling in microglia alters glycolytic capacity, we performed a glycolytic stress test on these cells using a Seahorse Analyzer. To our surprise, we found that glycolysis in the IR KD microglia was significantly increased (*P* < 0.01, Figures 4G and 4H), as was glycolytic capacity (*P* < 0.05, Figure 4I). In addition, non-glycolytic acidification (*P* < 0.01, Figure S8G), but not glycolytic reserve, was increased (Figure S8H). Thus, loss of IR in microglial cells leads to increased glycolysis and metabolic reprogramming.

### Loss of microglia IR signaling causes microglia activation and exaggerates AD pathology in 5xFAD mice

In addition to changes in mitochondrial metabolism, RiboTag profiling showed that IR deletion from microglia led to activation of innate immune pathways, including a more than 3-fold increased the levels of microglial signature genes, including *Tmem119*, *Hexb*, *Tyrobp*, *Itgam*, *Trem2*, and *Cd33* (Figures S9A and 9B). As shown above, microglia IRKO also resulted in increases in multiple genes in the complement pathway, including all three genes comprising the 18 polypeptide chains of C1q, namely *C1qa*, *C1qb*, and *C1qc* (Figure 3D). Recent studies have shown that activation of the microglial complement pathway can mediate loss of synapses and impairment in phagocytosis, leading to accelerated development of neurodegenerative diseases, such as Huntington’s disease and AD (Dejanovic et al., 2022; Wilton et al., 2023). It is also known that C1q, the initiator of the classical pathway of the complement system, is activated in AD, and this associated with the production and deposition of Aβ and phosphorylated tau in Aβ plaques and neurofibrillary tangles (Afagh et al., 1996; Dejanovic *et al*., 2022).

To test the hypothesis that activation of microglia induced by IR deletion could lead to altered Aβ uptake and aggravate the deposition of Aβ plaque during AD progression, we bred MG-IRKO mice with 5xFAD mice and assessed how loss of microglia IR signaling impacts on AD pathogenesis. Using a SHIELD-based CLARITY approach (Chen *et al*., 2023), we performed whole-brain mapping of Iba1, a pan microglia marker and marker of microglial activation (Ito et al., 1998; Walker and Lue, 2015). This found that compared to brains of mice carrying the 5xFAD transgenes only, brains of MG^IRKO^/5xFAD mice exhibited significantly higher levels of Iba1^+^ signals across multiple brain areas (Figures 5A and 5B), including the cerebral cortex, hippocampus, and hypothalamus, indicative of widespread neuroinflammation. Focusing on subregions in the prefrontal cortex and hippocampus involved in memory storage and retrieval, we found that both the dorsal and ventral hippocampus and entorhinal area had highest levels of neuroinflammation, as indicated by Iba1^+^ signals (Figures S9C and 9D). When compared to the 5xFAD control, the postsubiculum showed the greatest increase of Iba1^+^ signals (*P* < 0.001, Figure S9C), followed by the central amygdala. In the prefrontal cortex, the prelimbic area had the greatest increase of Iba1^+^ signals compared to the control (*P* < 0.01, Figure S9D). Thus, loss of microglia IR signaling leads to widespread of neuroinflammation across the whole-brain, potentially contributing to the AD pathogenesis.

**Figure 5.**
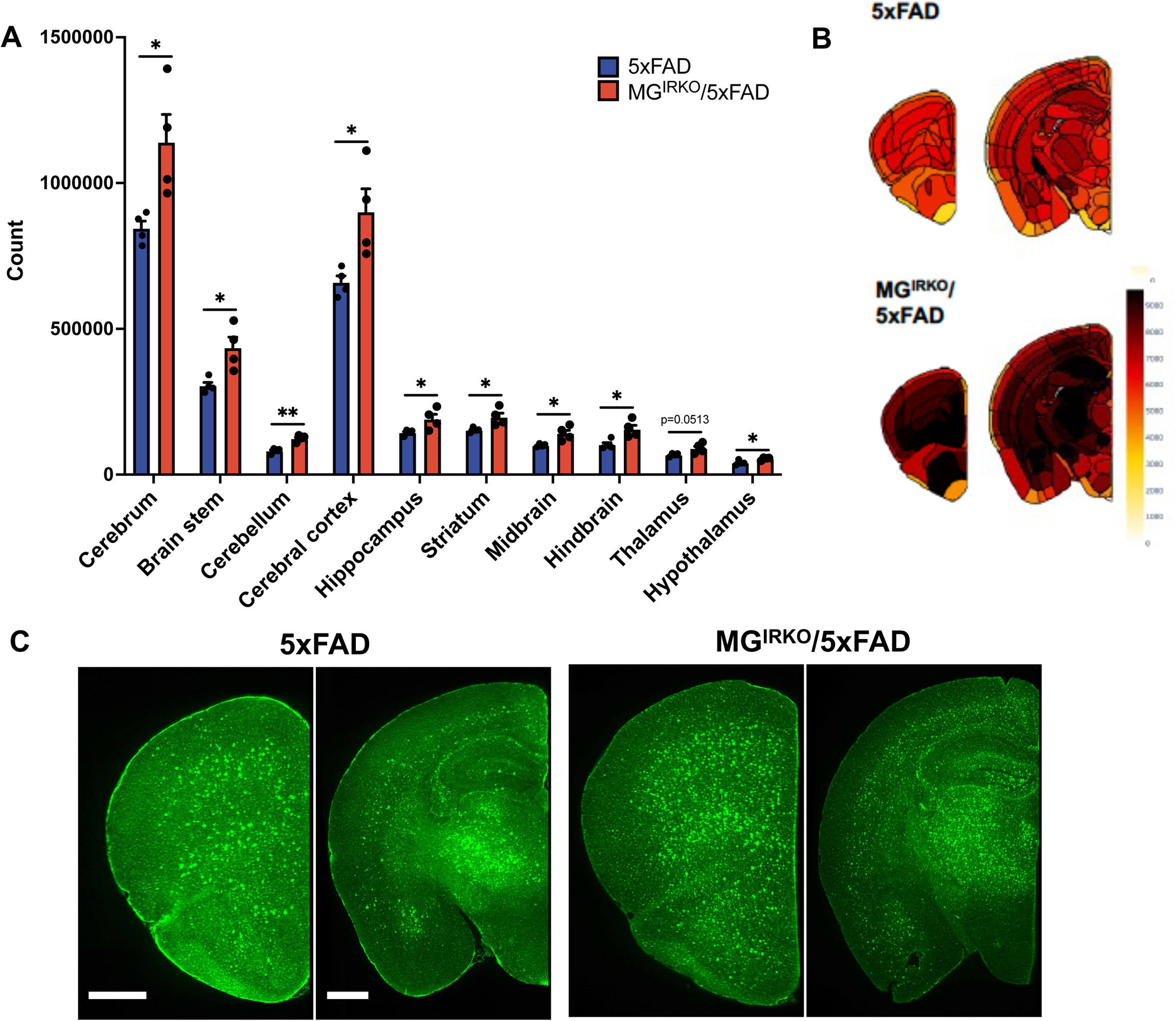
Loss of microglia IR signaling causes microglia activation and impairs Aβ uptake. (A-B) SHIELD-based whole-brain CLARITY analysis of IBA1 expression across the brain from MG^IRKO^/5xFAD mice compared to 5xFAD mice. N = 4. Bar graph (C) and representative heatmap graphs (B) showing expression levels of IBA1 in selective brain regions. Data are mean ± SEM. \**P* < 0.05, \*\**P* < 0.01, unpaired *t*-test. (C) representative fluorescent images of IBA1 expression in various brain regions. Scale bar, 100 µm.

In AD, it is known that the deposition of Aβ triggers activation of glial cells, including astrocytes and microglia, which initially try to protect against this process (Hansen et al., 2018). However, as AD advances, chronic activation of microglia leads to a change from a protective phenotype to a pro-inflammatory phenotype, resulting in accumulation of inflammation hallmarks in the brain, and this further exacerbates Aβ deposition and its spread throughout the brain (Leng and Edison, 2021). To test whether the increase in neuroinflammation due to microglial IR knockout was associated with an increased Aβ deposition, we performed immunohistochemical staining of the amyloid protein. Again, compared to 5xFAD mice, MG^IRKO^/5xFAD mice had increased Aβ deposition throughout the brain (Figure S10A), particularly in the subiculum (Figures 6A and 6B), a region of the brain with the earliest Aβ deposition (Gail Canter et al., 2019). Isolation of soluble and insoluble fractions of the brain hemispheres (Figure S10B) followed by ELISA assay revealed that brains from MG^IRKO^/5xFAD mice had a nearly 2-fold increase in deposition of insoluble Aβ_42_ (*P* < 0.01), but no increase in soluble Aβ_42_, compared with the 5xFAD mice (Figure 6C). By contrast, Aβ_40_ levels were much lower than levels of Aβ_42_ in all samples, and MG^IRKO^/5xFAD mice had significantly increased levels in the soluble fraction, but no change in the insoluble fraction (Figure S10C). Consistent with the brain CLARITY results showing widespread neuroinflammation, we found that levels of pro-inflammatory cytokine IFN-γ were increased ∼30% (*P* < 0.001) in the soluble fractions from MG^IRKO^/5xFAD brain (Figure 6D). This occurred with no change in the levels of IL-4 (Figure S10D). These data suggest that loss of insulin signaling in microglia exaggerates the neuroinflammation associated with AD pathogenesis.

**Figure 6.**
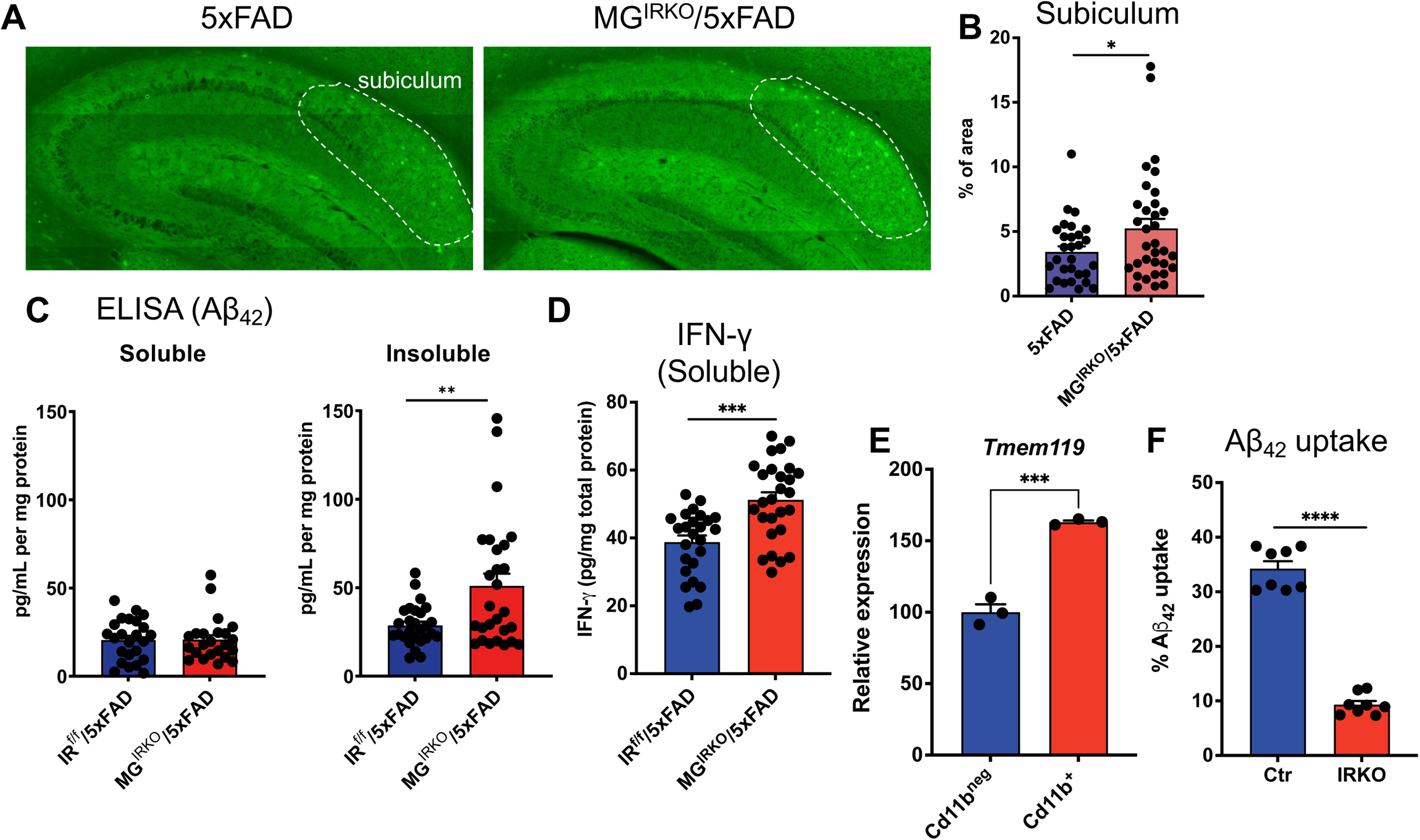
Loss of microglia IR signaling exaggerates AD pathology. (A) Representative fluorescent images of β-amyloid deposition in the subiculum from 6-month male MG^IRKO^/5xFAD mice compared to 5xFAD mice. (B) % of area in the subiculum from 6-month male MG^IRKO^/5xFAD mice compared to 5xFAD mice. N = 4 per group. (C) Aβ_42_ levels in soluble and insoluble fractions from brain homogenates of 6-month 5xFAD and MG^IRKO^/5xFAD mice; N = 27 per group. (D) ELISA for IFN-γ levels in soluble fraction from brain homogenates of 6-month 5xFAD and MG^IRKO^/5xFAD mice; N = 27 per group. (E) Relative expression of mRNA of *Tmem119* in microglia positive (Cd11b^+^ cells) and microglia negative cells (CD11b^−^ cells) isolated from male adult IR^f/f^ mice. N = 3 mice per group. unpaired *t*-test. (F) FACS analysis of uptake of Aβ_42_ in isolated microglia cells with IRKO. N = 8 per group. Data are mean ± SEM. \**P* < 0.05, \*\*\**P* < 0.001, \*\*\*\**P* < 0.0001, unpaired *t*-test.

Astrocytes and microglia are the major CNS phagocytes, and previous studies have shown that microglia are responsible for clearing necrotic tissue and cell debris, as well as phagocytosing Aβ deposits (Wang et al., 2021). We have previously demonstrated that insulin signaling in astrocytes can regulate uptake of Aβ (Chen *et al*., 2023), but whether insulin signaling in microglia regulates this function in unknown. To assess this, we isolated microglia from neonatal IR^f/f^ mice using Percoll gradient centrifugation followed by selection using anti-CD11b microbeads (Buenaventura et al., 2022). The resultant Cd11b^+^ cells showed enrichment of microglia marker genes, including *Tmem119*, *Tyrobp*, *Mef2c*, and *Cstd*, as compared to the non-microglial cells (CD11b^−^ cells), while *Clu*, a gene mainly expressed by astrocytes, was depleted in the Cd11b^+^ cells (Figure 6E and Figure S10E). We then infected the purified microglia with an adenovirus encoding a Cre:GFP fusion protein (or a control adenovirus) to induce deletion of the *Insr* gene. This resulted in a >60% decrease in *Insr* mRNA (Figure S10F), with no change in the insulin-like growth factor 1 receptor gene (*Igf1r*) (Figure S10G). When the control and IR knockdown microglia were incubated with fluorescently labeled Aβ_42_^HL-555^ and subjected to flow cytometry to measure Aβ uptake, the results show that compared to the control, microglia with IR KO exhibited an ∼40% decrease in uptake of Aβ peptide *in vitro* (Figure 6F). Thus, loss of IR signaling in microglia impairs uptake of Aβ.

## Discussion

Although the brain has traditionally been considered an insulin-insensitive tissue, multiple studies have shown that insulin signaling in brain can have effects on peripheral energy metabolism, as well as control of mood and cognition [reviewed in (Chen *et al*., 2022) and (Agrawal et al., 2021). Thus, brain samples from AD patients show altered insulin signaling (Gabbouj et al., 2019; Talbot *et al*., 2012), and mice with knockout of the insulin receptor in brain neuronal cells exhibit depressive-like behaviors and an exaggeration of Alzheimer’s phenotypes (Chen *et al*., 2023; Corraliza-Gomez et al., 2023; Jang et al., 2024). Likewise, loss of insulin signaling in astrocytes impairs cellular metabolism and Aβ uptake, and leads to the accelerated pathogenesis of AD (Cai *et al*., 2018; Chen *et al*., 2023). However, whether insulin signaling in microglia, the primary brain-resident immune cells, plays a role in brain function and AD pathogenesis remains unknown.

To answer these questions, we have established both cellular and mouse models in which insulin receptors have been selectively deleted in microglia (MG-IRKO). We find that, at the cellular level, loss of IR signaling in microglia results in metabolic reprograming with increased glycolysis and impaired uptake of Aβ peptide *in vitro*. *In vivo*, loss of microglial insulin signaling has minimal effects on systemic glucose metabolism or whole-body insulin sensitivity, but has significant effects on microglial gene expression, mood and social recognition behaviors, and progressive pathogenesis of AD. Thus, using RiboTag profiling to assess the role of insulin on gene expression in microglia, we demonstrate that loss of microglial insulin signaling leads to suppression of cellular metabolism, including fatty acid and branched chain amino acid metabolism and the TCA cycle, and activation of innate immune response pathways, including genes associated with phagocytosis, chemokine and cytokine pathways. As a result, mice with microglia IRKO exhibit increased depressive-like and stress-coping behaviors and altered social interaction. Furthermore, when the MG^IRKO^ mice are bred with the 5xFAD mouse model of AD, loss of insulin signaling in microglia exacerbates AD neuropathology. Thus, our findings demonstrate a critical role of insulin signaling in microglia to regulate cellular metabolism, the neuroinflammatory response and Aβ uptake leading to altered mood and social interactions and accentuating neurodegenerative diseases like AD.

Previous studies have shown that microglia and the brain immune system play key roles in modulation of mood, cognition, and behavior. Autistic behaviors can by induced by disrupting microglial-mediated synaptic pruning (Zhan et al., 2014), a primary function of microglia during brain development. The immune pathway can also play a role in behavior. Thus, mice deficient in adaptive immunity exhibit social deficit behaviors, mediated in part by proinflammatory cytokines, including IFN-γ (Filiano et al., 2016). Under pathological conditions, such as in multiple sclerosis and AD, microglia can be primed by IFN-γ to have an exaggerated immune response and shift to an activated state (Ta et al., 2019). Using a combination of cell specific knockout of IR and a RiboTag approach to assess gene expression, we show that loss of insulin signaling in microglia not only reduces expression of genes involved in metabolism, but exaggerates expression of genes in innate immune pathways, including both IFN-γ and its receptors. Furthermore, when mice lacking IR in microglial are crossed with the AD mouse model of 5xFAD, the resultant mice exhibit a higher degree of neuroinflammation, as demonstrated by increased expression of Iba1 and increased secretion of IFN-γ, and increased deposition of Aβ, especially in brain areas associated with cognition. Thus, our data point to insulin signaling as a key regulator of the immune response by microglia and their role in normal brain physiology.

Neuroinflammation has been well-recognized as a key pathophysiological feature of AD. In AD, the deposition of Aβ is an early key event. This triggers a microglial response as part of a neuroprotective role against AD development (Shi and Holtzman, 2018). However, as AD advances, chronic microglia activation causes reactive microglia to shift from this protective response to a more pro-inflammatory phenotype, resulting in accumulation of hallmarks of inflammation in the brain. This exaggerated inflammatory environment accelerates the formation of Aβ plaques from its initial seeding stage to a more widespread stage (Leng and Edison, 2021; Liu et al., 2023). Recent studies have identified Trem2 as a metabolic checkpoint for microglia function. Exclusively expressed in microglia, genetic mutation in *Trem2* can cause several-fold increased risks to develop AD (Keren-Shaul *et al*., 2017), and *Trem2* expression is increased 3-fold in MG^IRKO^ mice. Thus, insulin action controlling the expression of this and other immunoregulatory genes can play a major role in acceleration of the AD phenotypes.

Cellular metabolism in microglia also plays a critical role in immune surveillance (Bernier et al., 2020) contributing to the pathogenesis of AD (Baik et al., 2019). Consistent with this notion, our RiboTag profiling reveals a major effect of microglia IR KO on gene expression in pathways involved in cellular metabolism, including mitochondrial metabolism and regulation of autophagy/mitophagy. We show that in microglia cells, loss of insulin signaling reduces overall mitochondrial metabolism, especially through the TCA cycle, but facilitates glycolysis, indicative of metabolic reprogramming. This is consistent with the fact that, as the brain-resident macrophages, microglia are capable of switching from oxidative phosphorylation to glycolysis, a phenomenon well-characterized in tumor cells termed the Warburg effect (Liberti and Locasale, 2016). This allows for a rapid production of ATP so that microglia are prepared for activation. Interestingly, several recent studies have shown that in neurons derived *in vitro* from AD patient fibroblasts, there is also a metabolic switch toward glycolysis, and this is an important mechanism in promoting neuronal cell fate loss (Traxler et al., 2022). On the other hand, accumulation of Aβ itself can trigger microglia activation through metabolic reprogramming to glycolysis (Baik *et al*., 2019). Consistent with these findings, microglia-specific deletion of *Pkm2*, a gene encoding pyruvate kinase M2 (PKM2), attenuates microglial neuroinflammation and accumulation of Aβ (Pan et al., 2022), suggesting that reversing aberrant glycolysis in microglia may be a strategy to restore microglia function. Coupled with data demonstrating a key regulatory role of insulin signaling in microglia metabolism and immune regulation, these findings indicate that energy metabolism in microglia plays a critical role in mediating its immune responses upon pathological stimuli, such as accumulation of Aβ plaques, and that IR signaling is an important regulator of this process.

Deficits in cellular metabolism in microglia are also associated with impaired phagocytosis (Paolicelli *et al*., 2022). Phagocytosis is a core function of microglia, and alterations in this function can have profound impacts in pathogenesis of disease, including AD (Querfurth and LaFerla, 2010). Here we show that loss of insulin signaling in microglia causes a decrease in Aβ uptake, and in 5xFAD mice prone to AD, this loss of phagocytic potential results in accumulation of extracellular Aβ in the brain. During early stages of Aβ deposition, microglia form the first line of defense by engulfing and degrading Aβ (Baik *et al*., 2019; Querfurth and LaFerla, 2010). As microglia become reactive, more IFN-γ is secreted. This not only induces activation of the other cell types in the brain, but further impairs phagocytic activity of microglia. In line with our data, several studies have demonstrated a role of intranasal insulin on microglial responses. For example, in the 3xTg mouse model of AD, intranasal insulin administration has been shown to reduce Aβ levels and microglia activation (Chen *et al*., 2014). Likewise, in a senescence-accelerated mouse strain (SAMP8), intranasal insulin delivery shows beneficial effects on expression of inflammation-related genes in the hippocampus (Rhea et al., 2019), and in aged Fischer 344 rats, intranasal insulin administration has been shown to alter hippocampal gene expression in pathways associated with inflammation (Frazier *et al*., 2020). On the other hand, mice with genetic deficiency in adaptive immunity (SCID mice) exhibit social deficits, and this has been attributed to increased release of IFN-γ by T-cells in the meningeal compartment (Filiano *et al*., 2016).

In summary, the present study demonstrates a critical role of microglial insulin signaling in the regulation of microglial metabolism, neuroinflammation, cytokine secretion, and Aβ uptake, both *in vitro* and *in vivo*, and in the latter, this contributes to accelerated amyloid pathology in a mouse model of AD. This fits well with studies of cerebrospinal fluid (CSF) proteomics which have demonstrated the involvement of microglia activation and energy metabolism in humans with AD (Johnson et al., 2020), as well as a recent CSF proteomics study which has identified several AD subtypes, including one subtype that is characterized by a core signature of innate immune activation (Tijms et al., 2024). Our findings point to the importance of understanding a cell type-specific regulation of insulin action and insulin resistance in brain homeostasis and disease pathogenesis. These insights will help identify the cellular and molecular mechanisms underlying the comorbidity of T2D and brain disorders, thus bringing the potential for better therapeutics for patients with these comorbid conditions.

### Limitations of Study

While our study shows a key role of microglial insulin signaling in brain functions, we have not investigated the roles of IGF-1 receptors (IGF1R) signaling in microglia. Given that insulin can also bind to the IGF1R, a related tyrosine kinase receptor that is normally activated by IGF-1 and IGF-2, and that both IR and IGF1R are expressed on microglia (Martinez-Rachadell et al., 2019), it is possible that the IGF1R signaling might exert a similar, but distinct, role to that of the IR signaling in microglia. Further, the genetically modified mice used in our study are mice with KO of IR in TMEM119-expessing cells. While this includes microglia in the brain parenchyma (Kaiser and Feng, 2019), it is possible that there are minor levels of expression in other cell types, such as meningeal T cells mentioned above. In other studies, microglia have been targeted by using the *Cx3cr1* promoter-driven Cre lines (Haimon et al., 2018; Sahasrabuddhe and Ghosh, 2022; Zhang et al., 2018; Zhao et al., 2018), as well as *Hexb^CreERT2^* (Masuda et al., 2020), *P2ry12^CreER^*(McKinsey et al., 2020), and *Sall1^CreER^* mouse lines (Buttgereit et al., 2016). While each of these can target populations of microglia, at this time, the TMEM119 promoter seems to offer the greatest specificity. Another limitation worth noting is that none of these approaches modify microglia insulin signaling in a brain region-specific manner, and to the extent that microglia exhibit region-specific signatures (Tan et al., 2020), the heterogeneity of microglia physiology at a subregion level would be a topic of interest for future experimental studies.

## Supporting information

Supplemental infomation

## Acknowledgments

This work was supported by NIH grants R01 DK031036 (to C.R.K.), in part by NIH training grant T32 DK007260 (to W.C.) and in part by the Steno North American Fellowship (NNF23OC0087108) awarded by the Novo Nordisk Foundation (to W.C.). The Joslin Diabetes Center DRC Advanced Microscopy Core, Flow Cytometry Core, and Animal Physiology Core (P30 DK036836) provided important help. We thank LifeCanvas Inc. for providing important help on whole brain clearing, imaging, and analysis. We thank the support of the HMS Neurobiology Imaging Facility (supported by NINDS Core Center Grant P30NS072030). We thank Dr. Hongbin Yang (Zhejiang University) for critical reading of the manuscript. We thank Christopher Cahill and Emily Stephens for their technical assistance and thank all the Kahn lab members for their helpful discussion. Schematics are made with Biorender.com.

## REFERENCES

Afagh, A., Cummings, B.J., Cribbs, D.H., Cotman, C.W., and Tenner, A.J. (1996). Localization and cell association of C1q in Alzheimer’s disease brain. Exp Neurol 138, 22–32. 10.1006/exnr.1996.0043.

Agrawal, R., Reno, C.M., Sharma, S., Christensen, C., Huang, Y., and Fisher, S.J. (2021). Insulin action in the brain regulates both central and peripheral functions. Am J Physiol Endocrinol Metab 321, E156–E163. 10.1152/ajpendo.00642.2020.

Alers, S., Loffler, A.S., Wesselborg, S., and Stork, B. (2012). Role of AMPK-mTOR-Ulk1/2 in the regulation of autophagy: cross talk, shortcuts, and feedbacks. Molecular and cellular biology 32, 2–11. 10.1128/MCB.06159-11.

Arnold, S.E., Arvanitakis, Z., Macauley-Rambach, S.L., Koenig, A.M., Wang, H.Y., Ahima, R.S., Craft, S., Gandy, S., Buettner, C., Stoeckel, L.E., et al. (2018). Brain insulin resistance in type 2 diabetes and Alzheimer disease: concepts and conundrums. Nature reviews. Neurology 14, 168–181. 10.1038/nrneurol.2017.185.

Baik, S.H., Kang, S., Lee, W., Choi, H., Chung, S., Kim, J.I., and Mook-Jung, I. (2019). A Breakdown in Metabolic Reprogramming Causes Microglia Dysfunction in Alzheimer’s Disease. Cell Metab 30, 493–507 e496. 10.1016/j.cmet.2019.06.005.

Barbagallo, M., and Dominguez, L.J. (2014). Type 2 diabetes mellitus and Alzheimer’s disease. World J Diabetes 5, 889–893. 10.4239/wjd.v5.i6.889.

Bennett, M.L., Bennett, F.C., Liddelow, S.A., Ajami, B., Zamanian, J.L., Fernhoff, N.B., Mulinyawe, S.B., Bohlen, C.J., Adil, A., Tucker, A., et al. (2016). New tools for studying microglia in the mouse and human CNS. Proceedings of the National Academy of Sciences of the United States of America 113, E1738–1746. 10.1073/pnas.1525528113.

Bernier, L.P., York, E.M., Kamyabi, A., Choi, H.B., Weilinger, N.L., and MacVicar, B.A. (2020). Microglial metabolic flexibility supports immune surveillance of the brain parenchyma. Nature communications 11, 1559. 10.1038/s41467-020-15267-z.

Bruning, J.C., Gautam, D., Burks, D.J., Gillette, J., Schubert, M., Orban, P.C., Klein, R., Krone, W., Muller-Wieland, D., and Kahn, C.R. (2000). Role of brain insulin receptor in control of body weight and reproduction. Science 289, 2122–2125. 10.1126/science.289.5487.2122.

Bruning, J.C., Michael, M.D., Winnay, J.N., Hayashi, T., Horsch, D., Accili, D., Goodyear, L.J., and Kahn, C.R. (1998). A muscle-specific insulin receptor knockout exhibits features of the metabolic syndrome of NIDDM without altering glucose tolerance. Mol Cell 2, 559–569.

Buenaventura, R.G., Harvey, A.C., Burns, M.P., and Main, B.S. (2022). Sequential Isolation of Microglia and Astrocytes from Young and Aged Adult Mouse Brains for Downstream Transcriptomic Analysis. Methods Protoc 5. 10.3390/mps5050077.

Buttgereit, A., Lelios, I., Yu, X., Vrohlings, M., Krakoski, N.R., Gautier, E.L., Nishinakamura, R., Becher, B., and Greter, M. (2016). Sall1 is a transcriptional regulator defining microglia identity and function. Nature immunology 17, 1397–1406. 10.1038/ni.3585.

Cai, W., Xue, C., Sakaguchi, M., Konishi, M., Shirazian, A., Ferris, H.A., Li, M.E., Yu, R., Kleinridders, A., Pothos, E.N., and Kahn, C.R. (2018). Insulin regulates astrocyte gliotransmission and modulates behavior. The Journal of clinical investigation 128, 2914–2926. 10.1172/JCI99366.

Cai, W., Zhang, X., Batista, T.M., Garcia-Martin, R., Softic, S., Wang, G., Ramirez, A.K., Konishi, M., O’Neill, B.T., Kim, J.H., et al. (2021). Peripheral Insulin Regulates a Broad Network of Gene Expression in the Hypothalamus, Hippocampus and Nucleus Accumbens. Diabetes. 10.2337/db20-1119.

Castagne, V., Moser, P., Roux, S., and Porsolt, R.D. (2011). Rodent models of depression: forced swim and tail suspension behavioral despair tests in rats and mice. Curr Protoc Neurosci Chapter 8, Unit 8 10A. 10.1002/0471142301.ns0810as55.

Chen, W., Cai, W., Hoover, B., and Kahn, C.R. (2022). Insulin action in the brain: cell types, circuits, and diseases. Trends in neurosciences 45, 384–400. 10.1016/j.tins.2022.03.001.

Chen, W., Huang, Q., Lazdon, E.K., Gomes, A., Wong, M., Stephens, E., Royal, T.G., Frenkel, D., Cai, W., and Kahn, C.R. (2023). Loss of insulin signaling in astrocytes exacerbates Alzheimer-like phenotypes in a 5xFAD mouse model. Proc Natl Acad Sci U S A 120, e2220684120. 10.1073/pnas.2220684120.

Chen, Y., Zhao, Y., Dai, C.L., Liang, Z., Run, X., Iqbal, K., Liu, F., and Gong, C.X. (2014). Intranasal insulin restores insulin signaling, increases synaptic proteins, and reduces Abeta level and microglia activation in the brains of 3xTg-AD mice. Exp Neurol 261, 610–619. 10.1016/j.expneurol.2014.06.004.

Chhimpa, N., Singh, N., Puri, N., and Kayath, H.P. (2023). The Novel Role of Mitochondrial Citrate Synthase and Citrate in the Pathophysiology of Alzheimer’s Disease. J Alzheimers Dis 94, S453–S472. 10.3233/JAD-220514.

Corraliza-Gomez, M., Bermejo, T., Lilue, J., Rodriguez-Iglesias, N., Valero, J., Cozar-Castellano, I., Arranz, E., Sanchez, D., and Ganfornina, M.D. (2023). Insulin-degrading enzyme (IDE) as a modulator of microglial phenotypes in the context of Alzheimer’s disease and brain aging. Journal of neuroinflammation 20, 233. 10.1186/s12974-023-02914-7.

Dejanovic, B., Wu, T., Tsai, M.C., Graykowski, D., Gandham, V.D., Rose, C.M., Bakalarski, C.E., Ngu, H., Wang, Y., Pandey, S., et al. (2022). Complement C1q-dependent excitatory and inhibitory synapse elimination by astrocytes and microglia in Alzheimer’s disease mouse models. Nat Aging 2, 837–850. 10.1038/s43587-022-00281-1.

Filiano, A.J., Xu, Y., Tustison, N.J., Marsh, R.L., Baker, W., Smirnov, I., Overall, C.C., Gadani, S.P., Turner, S.D., Weng, Z., et al. (2016). Unexpected role of interferon-gamma in regulating neuronal connectivity and social behaviour. Nature 535, 425–429. 10.1038/nature18626.

Frazier, H.N., Ghoweri, A.O., Sudkamp, E., Johnson, E.S., Anderson, K.L., Fox, G., Vatthanaphone, K., Xia, M., Lin, R.L., Hargis-Staggs, K.E., et al. (2020). Long-Term Intranasal Insulin Aspart: A Profile of Gene Expression, Memory, and Insulin Receptors in Aged F344 Rats. J Gerontol A Biol Sci Med Sci 75, 1021–1030. 10.1093/gerona/glz105.

Frendo-Cumbo, S., Tokarz, V.L., Bilan, P.J., Brumell, J.H., and Klip, A. (2021). Communication Between Autophagy and Insulin Action: At the Crux of Insulin Action-Insulin Resistance? Front Cell Dev Biol 9, 708431. 10.3389/fcell.2021.708431.

Frolich, L., Blum-Degen, D., Bernstein, H.G., Engelsberger, S., Humrich, J., Laufer, S., Muschner, D., Thalheimer, A., Turk, A., Hoyer, S., et al. (1998). Brain insulin and insulin receptors in aging and sporadic Alzheimer’s disease. J Neural Transm (Vienna) 105, 423–438. 10.1007/s007020050068.

Gabbouj, S., Ryhanen, S., Marttinen, M., Wittrahm, R., Takalo, M., Kemppainen, S., Martiskainen, H., Tanila, H., Haapasalo, A., Hiltunen, M., and Natunen, T. (2019). Altered Insulin Signaling in Alzheimer’s Disease Brain - Special Emphasis on PI3K-Akt Pathway. Frontiers in neuroscience 13, 629. 10.3389/fnins.2019.00629.

Gail Canter, R., Huang, W.C., Choi, H., Wang, J., Ashley Watson, L., Yao, C.G., Abdurrob, F., Bousleiman, S.M., Young, J.Z., Bennett, D.A., et al. (2019). 3D mapping reveals network-specific amyloid progression and subcortical susceptibility in mice. Commun Biol 2, 360. 10.1038/s42003-019-0599-8.

Geffken, S.J., Moon, S., Smith, C.O., Tang, S., Lee, H.H., Lewis, K., Wong, C.W., Huang, Y., Huang, Q., Zhao, Y.T., and Cai, W. (2022). Insulin and IGF-1 elicit robust transcriptional regulation to modulate autophagy in astrocytes. Mol Metab 66, 101647. 10.1016/j.molmet.2022.101647.

Haimon, Z., Volaski, A., Orthgiess, J., Boura-Halfon, S., Varol, D., Shemer, A., Yona, S., Zuckerman, B., David, E., Chappell-Maor, L., et al. (2018). Re-evaluating microglia expression profiles using RiboTag and cell isolation strategies. Nature immunology 19, 636–644. 10.1038/s41590-018-0110-6.

Hansen, D.V., Hanson, J.E., and Sheng, M. (2018). Microglia in Alzheimer’s disease. The Journal of cell biology 217, 459–472. 10.1083/jcb.201709069.

Ito, D., Imai, Y., Ohsawa, K., Nakajima, K., Fukuuchi, Y., and Kohsaka, S. (1998). Microglia-specific localisation of a novel calcium binding protein, Iba1. Brain research. Molecular brain research 57, 1–9. 10.1016/s0169-328x(98)00040-0.

Jang, Y.J., Choi, M.G., Yoo, B.J., Lee, K.J., Jung, W.B., Kim, S.G., and Park, S.A. (2024). Interaction Between a High-Fat Diet and Tau Pathology in Mice: Implications for Alzheimer’s Disease. J Alzheimers Dis 97, 485–506. 10.3233/JAD-230927.

Jiang, Z.H., Peng, J., Yang, H.L., Fu, X.L., Wang, J.Z., Liu, L., Jiang, J.N., Tan, Y.F., and Ge, Z.J. (2017). Upregulation and biological function of transmembrane protein 119 in osteosarcoma. Exp Mol Med 49, e329. 10.1038/emm.2017.41.

Johnson, E.C.B., Dammer, E.B., Duong, D.M., Ping, L., Zhou, M., Yin, L., Higginbotham, L.A., Guajardo, A., White, B., Troncoso, J.C., et al. (2020). Large-scale proteomic analysis of Alzheimer’s disease brain and cerebrospinal fluid reveals early changes in energy metabolism associated with microglia and astrocyte activation. Nat Med 26, 769–780. 10.1038/s41591-020-0815-6.

Kaiser, T., and Feng, G. (2019). Tmem119-EGFP and Tmem119-CreERT2 Transgenic Mice for Labeling and Manipulating Microglia. eNeuro 6. 10.1523/ENEURO.0448-18.2019.

Kellar, D., and Craft, S. (2020). Brain insulin resistance in Alzheimer’s disease and related disorders: mechanisms and therapeutic approaches. Lancet neurology 19, 758–766. 10.1016/S1474-4422(20)30231-3.

Keren-Shaul, H., Spinrad, A., Weiner, A., Matcovitch-Natan, O., Dvir-Szternfeld, R., Ulland, T.K., David, E., Baruch, K., Lara-Astaiso, D., Toth, B., et al. (2017). A Unique Microglia Type Associated with Restricting Development of Alzheimer’s Disease. Cell 169, 1276–1290 e1217. 10.1016/j.cell.2017.05.018.

Kim, J., Kundu, M., Viollet, B., and Guan, K.L. (2011). AMPK and mTOR regulate autophagy through direct phosphorylation of Ulk1. Nature cell biology 13, 132–141. 10.1038/ncb2152.

Kim, J.D., Yoon, N.A., Jin, S., and Diano, S. (2019). Microglial UCP2 Mediates Inflammation and Obesity Induced by High-Fat Feeding. Cell metabolism 30, 952–962 e955. 10.1016/j.cmet.2019.08.010.

Leng, F., and Edison, P. (2021). Neuroinflammation and microglial activation in Alzheimer disease: where do we go from here? Nature reviews. Neurology 17, 157–172. 10.1038/s41582-020-00435-y.

Liberti, M.V., and Locasale, J.W. (2016). The Warburg Effect: How Does it Benefit Cancer Cells? Trends Biochem Sci 41, 211–218. 10.1016/j.tibs.2015.12.001.

Liu, C.C., Wang, N., Chen, Y., Inoue, Y., Shue, F., Ren, Y., Wang, M., Qiao, W., Ikezu, T.C., Li, Z., et al. (2023). Cell-autonomous effects of APOE4 in restricting microglial response in brain homeostasis and Alzheimer’s disease. Nature immunology 24, 1854–1866. 10.1038/s41590-023-01640-9.

Martinez-Rachadell, L., Aguilera, A., Perez-Domper, P., Pignatelli, J., Fernandez, A.M., and Torres-Aleman, I. (2019). Cell-specific expression of insulin/insulin-like growth factor-I receptor hybrids in the mouse brain. Growth Horm IGF Res 45, 25–30. 10.1016/j.ghir.2019.02.003.

Masuda, T., Amann, L., Sankowski, R., Staszewski, O., Lenz, M., P, D.E., Snaidero, N., Costa Jordao, M.J., Bottcher, C., Kierdorf, K., et al. (2020). Novel Hexb-based tools for studying microglia in the CNS. Nature immunology 21, 802–815. 10.1038/s41590-020-0707-4.

McKinsey, G.L., Lizama, C.O., Keown-Lang, A.E., Niu, A., Santander, N., Larpthaveesarp, A., Chee, E., Gonzalez, F.F., and Arnold, T.D. (2020). A new genetic strategy for targeting microglia in development and disease. Elife 9. 10.7554/eLife.54590.

Mordelt, A., and de Witte, L.D. (2023). Microglia-mediated synaptic pruning as a key deficit in neurodevelopmental disorders: Hype or hope? Current opinion in neurobiology 79, 102674. 10.1016/j.conb.2022.102674.

Nagamoto-Combs, K., Kulas, J., and Combs, C.K. (2014). A novel cell line from spontaneously immortalized murine microglia. Journal of neuroscience methods 233, 187–198. 10.1016/j.jneumeth.2014.05.021.

Orihuela, R., McPherson, C.A., and Harry, G.J. (2016). Microglial M1/M2 polarization and metabolic states. Br J Pharmacol 173, 649–665. 10.1111/bph.13139.

Pan, R.-Y., He, L., Zhang, J., Liu, X., Liao, Y., Gao, J., Liao, Y., Yan, Y., Li, Q., Zhou, X., et al. (2022). Positive feedback regulation of microglial glucose metabolism by histone H4 lysine 12 lactylation in Alzheimer&#x2019;s disease. Cell Metabolism 34, 634–648.e636. 10.1016/j.cmet.2022.02.013.

Paolicelli, R.C., Sierra, A., Stevens, B., Tremblay, M.E., Aguzzi, A., Ajami, B., Amit, I., Audinat, E., Bechmann, I., Bennett, M., et al. (2022). Microglia states and nomenclature: A field at its crossroads. Neuron 110, 3458–3483. 10.1016/j.neuron.2022.10.020.

Park, J.M., Lee, D.H., and Kim, D.H. (2023). Redefining the role of AMPK in autophagy and the energy stress response. Nature communications 14, 2994. 10.1038/s41467-023-38401-z.

Querfurth, H.W., and LaFerla, F.M. (2010). Alzheimer’s disease. The New England journal of medicine 362, 329–344. 10.1056/NEJMra0909142.

Rhea, E.M., Nirkhe, S., Nguyen, S., Pemberton, S., Bammler, T.K., Beyer, R., Niehoff, M.L., Morley, J.E., Farr, S.A., and Banks, W.A. (2019). Molecular Mechanisms of Intranasal Insulin in SAMP8 Mice. J Alzheimers Dis 71, 1361–1373. 10.3233/JAD-190707.

Roach, P.J. (2011). AMPK -> ULK1 -> autophagy. Molecular and cellular biology 31, 3082–3084. 10.1128/MCB.05565-11.

Ruan, C., Sun, L., Kroshilina, A., Beckers, L., De Jager, P., Bradshaw, E.M., Hasson, S.A., Yang, G., and Elyaman, W. (2020). A novel Tmem119-tdTomato reporter mouse model for studying microglia in the central nervous system. Brain Behav Immun 83, 180–191. 10.1016/j.bbi.2019.10.009.

Sahasrabuddhe, V., and Ghosh, H.S. (2022). Cx3Cr1-Cre induction leads to microglial activation and IFN-1 signaling caused by DNA damage in early postnatal brain. Cell Rep 38, 110252. 10.1016/j.celrep.2021.110252.

Sanz, E., Bean, J.C., Carey, D.P., Quintana, A., and McKnight, G.S. (2019). RiboTag: Ribosomal Tagging Strategy to Analyze Cell-Type-Specific mRNA Expression In Vivo. Curr Protoc Neurosci 88, e77. 10.1002/cpns.77.

Sanz, E., Yang, L., Su, T., Morris, D.R., McKnight, G.S., and Amieux, P.S. (2009). Cell-type-specific isolation of ribosome-associated mRNA from complex tissues. Proceedings of the National Academy of Sciences of the United States of America 106, 13939–13944. 10.1073/pnas.0907143106.

Satoh, J., Kino, Y., Asahina, N., Takitani, M., Miyoshi, J., Ishida, T., and Saito, Y. (2016). TMEM119 marks a subset of microglia in the human brain. Neuropathology 36, 39–49. 10.1111/neup.12235.

Schubert, M., Gautam, D., Surjo, D., Ueki, K., Baudler, S., Schubert, D., Kondo, T., Alber, J., Galldiks, N., Kustermann, E., et al. (2004). Role for neuronal insulin resistance in neurodegenerative diseases. Proc Natl Acad Sci U S A 101, 3100–3105. 10.1073/pnas.0308724101.

Sellgren, C.M., Gracias, J., Watmuff, B., Biag, J.D., Thanos, J.M., Whittredge, P.B., Fu, T., Worringer, K., Brown, H.E., Wang, J., et al. (2019). Increased synapse elimination by microglia in schizophrenia patient-derived models of synaptic pruning. Nature neuroscience 22, 374–385. 10.1038/s41593-018-0334-7.

Shi, Y., and Holtzman, D.M. (2018). Interplay between innate immunity and Alzheimer disease: APOE and TREM2 in the spotlight. Nat Rev Immunol 18, 759–772. 10.1038/s41577-018-0051-1.

Steinberg, G.R., and Carling, D. (2019). AMP-activated protein kinase: the current landscape for drug development. Nat Rev Drug Discov 18, 527–551. 10.1038/s41573-019-0019-2.

Ta, T.T., Dikmen, H.O., Schilling, S., Chausse, B., Lewen, A., Hollnagel, J.O., and Kann, O. (2019). Priming of microglia with IFN-gamma slows neuronal gamma oscillations in situ. Proceedings of the National Academy of Sciences of the United States of America 116, 4637–4642. 10.1073/pnas.1813562116.

Talbot, K., Wang, H.Y., Kazi, H., Han, L.Y., Bakshi, K.P., Stucky, A., Fuino, R.L., Kawaguchi, K.R., Samoyedny, A.J., Wilson, R.S., et al. (2012). Demonstrated brain insulin resistance in Alzheimer’s disease patients is associated with IGF-1 resistance, IRS-1 dysregulation, and cognitive decline. The Journal of clinical investigation 122, 1316–1338. 10.1172/JCI59903.

Tan, Y.L., Yuan, Y., and Tian, L. (2020). Microglial regional heterogeneity and its role in the brain. Mol Psychiatry 25, 351–367. 10.1038/s41380-019-0609-8.

Tanaka, K., Inoue, Y., Hendy, G.N., Canaff, L., Katagiri, T., Kitazawa, R., Komori, T., Sugimoto, T., Seino, S., and Kaji, H. (2012). Interaction of Tmem119 and the bone morphogenetic protein pathway in the commitment of myoblastic into osteoblastic cells. Bone 51, 158–167. 10.1016/j.bone.2012.04.017.

Tijms, B.M., Vromen, E.M., Mjaavatten, O., Holstege, H., Reus, L.M., van der Lee, S., Wesenhagen, K.E.J., Lorenzini, L., Vermunt, L., Venkatraghavan, V., et al. (2024). Cerebrospinal fluid proteomics in patients with Alzheimer’s disease reveals five molecular subtypes with distinct genetic risk profiles. Nat Aging 4, 33–47. 10.1038/s43587-023-00550-7.

Traxler, L., Herdy, J.R., Stefanoni, D., Eichhorner, S., Pelucchi, S., Szucs, A., Santagostino, A., Kim, Y., Agarwal, R.K., Schlachetzki, J.C.M., et al. (2022). Warburg-like metabolic transformation underlies neuronal degeneration in sporadic Alzheimer’s disease. Cell metabolism 34, 1248–1263 e1246. 10.1016/j.cmet.2022.07.014.

Walker, D.G., and Lue, L.F. (2015). Immune phenotypes of microglia in human neurodegenerative disease: challenges to detecting microglial polarization in human brains. Alzheimers Res Ther 7, 56. 10.1186/s13195-015-0139-9.

Wang, K., Li, J., Zhang, Y., Huang, Y., Chen, D., Shi, Z., Smith, A.D., Li, W., and Gao, Y. (2021). Central nervous system diseases related to pathological microglial phagocytosis. CNS Neurosci Ther 27, 528–539. 10.1111/cns.13619.

Wang, L., Pavlou, S., Du, X., Bhuckory, M., Xu, H., and Chen, M. (2019). Glucose transporter 1 critically controls microglial activation through facilitating glycolysis. Mol Neurodegener 14, 2. 10.1186/s13024-019-0305-9.

Wilton, D.K., Mastro, K., Heller, M.D., Gergits, F.W., Willing, C.R., Fahey, J.B., Frouin, A., Daggett, A., Gu, X., Kim, Y.A., et al. (2023). Microglia and complement mediate early corticostriatal synapse loss and cognitive dysfunction in Huntington’s disease. Nat Med. 10.1038/s41591-023-02566-3.

Wong, C.Y.J., Baldelli, A., Hoyos, C.M., Tietz, O., Ong, H.X., and Traini, D. (2024). Insulin Delivery to the Brain via the Nasal Route: Unraveling the Potential for Alzheimer’s Disease Therapy. Drug Deliv Transl Res 14, 1776–1793. 10.1007/s13346-024-01558-1.

Wu, J., Zhang, J., Chen, X., Wettschurack, K., Que, Z., Deming, B.A., Olivero-Acosta, M.I., Cui, N., Eaton, M., Zhao, Y., et al. (2024). Microglial over-pruning of synapses during development in autism-associated SCN2A-deficient mice and human cerebral organoids. Mol Psychiatry. 10.1038/s41380-024-02518-4.

Xu, X.J., Gauthier, M.S., Hess, D.T., Apovian, C.M., Cacicedo, J.M., Gokce, N., Farb, M., Valentine, R.J., and Ruderman, N.B. (2012). Insulin sensitive and resistant obesity in humans: AMPK activity, oxidative stress, and depot-specific changes in gene expression in adipose tissue. Journal of lipid research 53, 792–801. 10.1194/jlr.P022905.

Zhan, Y., Paolicelli, R.C., Sforazzini, F., Weinhard, L., Bolasco, G., Pagani, F., Vyssotski, A.L., Bifone, A., Gozzi, A., Ragozzino, D., and Gross, C.T. (2014). Deficient neuron-microglia signaling results in impaired functional brain connectivity and social behavior. Nature neuroscience 17, 400–406. 10.1038/nn.3641.

Zhang, B., Zou, J., Han, L., Beeler, B., Friedman, J.L., Griffin, E., Piao, Y.S., Rensing, N.R., and Wong, M. (2018). The specificity and role of microglia in epileptogenesis in mouse models of tuberous sclerosis complex. Epilepsia 59, 1796–1806. 10.1111/epi.14526.

Zhao, X., Liao, Y., Morgan, S., Mathur, R., Feustel, P., Mazurkiewicz, J., Qian, J., Chang, J., Mathern, G.W., Adamo, M.A., et al. (2018). Noninflammatory Changes of Microglia Are Sufficient to Cause Epilepsy. Cell Rep 22, 2080–2093. 10.1016/j.celrep.2018.02.004.

