## Supplemental infomation for "Loss of Insulin Signaling in Microglia Impairs Cellular Uptake of Aβ and Neuroinflammatory Response Exacerbating Alzheimer-like Neuropathology"

**This manuscript includes:**

Supplement Figures 1 to 10
STAR methods

### STAR METHODS

### KEY RESOURCES TABLE

| REAGENT or RESORCE | SOURCE | IDENTIFIER |
|---|---|---|
| **Antibodies** | | |
| Mouse Monoclonal anti-HA.11 (16B12) | BioLegend | Cat# MMS-101R, RRID:AB_291262 |
| Rat Monoclonal anti-CD11b (M1/70), Biotin | eBioscience | Cat# 13-0112-85, RRID:AB_466361 |
| Rabbit Polyclonal anti-GFAP | EMD Millipore | Cat# AB5804; RRID:AB_305124 |
| Rabbit Polyclonal anti-Iba1 | Fujifilm Wako | Cat# 27030; RRID:AB_2314667 |
| Rabbit Polyclonal anti-Iba1 | Fujifilm Wako | Cat# 019-19741; RRID:AB_839504 |
| Mouse Monoclonal anti-β-Amyloid (1-16) | BioLegend | Cat# 803001; RRID:AB_2564652 |
| Rabbit Polyclonal anti-β-Amyloid 1-42 (mOC64) | Abcam | Cat# ab201060, RRID:AB_2818982 |
| Rabbit Monoclonal anti-GAPDH (D16H11) | Cell Signaling Technology | Cat# 5174, RRID:AB_10622025 |
| Rabbit Monoclonal anti-phospho-AMPKα1 (Thr172) (40H9) | Cell Signaling Technology | Cat# 2535, RRID:AB_331250 |
| Rabbit Monoclonal anti-AMPKα1 (D5A2) | Cell Signaling Technology | Cat# 5831, RRID:AB_10622186 |
| Rabbit Monoclonal anti-phospho-mTOR (Ser2448) (D9C2) | Cell Signaling Technology | Cat# 5536, RRID:AB_10691552 |
| Rabbit Polyclonal anti-mTOR | Cell Signaling Technology | Cat# 2972, RRID:AB_330978 |
| Rabbit Monoclonal anti-phospho-ULK1 (Ser757) (D7O6U) | Cell Signaling Technology | Cat# 14202, RRID:AB_2665508 |
| Rabbit Monoclonal anti-ULK1 (D8H5) | Cell Signaling Technology | Cat# 8054, RRID:AB_11178668 |
| MOuse Monoclonal anti-Mitofusin-2 (6A8) | Abcam | Cat# ab56889, RRID:AB_2142629 |
| Rabbit Polyclonal anti-Citrate synthase | Abcam | Cat# ab96600, RRID:AB_10678258 |
| Total OXPHOS Rodent WB Antibody Cocktail | Abcam | Cat# ab110413, RRID:AB_2629281 |
| Rabbit Monoclonal anti-β-Actin (13E5) | Cell Signaling Technology | Cat# 4970, RRID:AB_2223172 |
| Rabbit Monoclonal anti-LC3A/B (D3U4C) | Cell Signaling Technology | Cat# 12741, RRID:AB_2617131 |
| Rabbit Monoclonal anti-SQSTM1/p62 | Cell Signaling Technology | Cat# 5114, RRID:AB_10624872 |
| Rabbit Polyclonal anti-Atg3 | Cell Signaling Technology | Cat# 3415, RRID:AB_2059244 |
| Rabbit Polyclonal anti-Atg5 | Cell Signaling Technology | Cat# 2630, RRID:AB_2062340 |
| Rabbit Polyclonal anti-Atg7 | Cell Signaling Technology | Cat# 2631, RRID:AB_2227783 |
| Rabbit Polyclonal anti-phospho-DRP1 (Ser637) | Cell Signaling Technology | Cat# 4867, RRID:AB_10622027 |
| Rabbit Polyclonal anti- phospho-DRP1 (Ser616) | Cell Signaling Technology | Cat# 3455, RRID:AB_2085352 |
| Rabbit Monoclonal anti-DRP1 (D8H5) | Cell Signaling Technology | Cat# 5391, RRID:AB_11178938 |
| Rabbit Polyclonal anti-FIS1 (FL-152) | Santa Cruz Biotechnology | Cat# sc-98900, RRID:AB_2246809 |
| Rabbit Polyclonal anti-Parkin | Cell Signaling Technology | Cat# 2132, RRID:AB_10693040 |
| Anti-Rat IgG | Jackson ImmunoResearch | Cat# 112-005-167; RRID:AB_2338101 |
| Anti-Mouse IgG, Alexa Fluor 647 | Invitrogen | Cat# A-21235; RRID:AB_2535804 |
| Anti-Rabbit IgG, Alexa Fluor 647 | Invitrogen | Cat# A-21244; RRID:AB_2535812 |
| Anti-Rabbit IgG, Alexa Fluor 647 | Invitrogen | Cat# A-21244; RRID:AB_2535812 |
| **Bacterial and virus strains** | | |
| $AAV_{2/9}$-CAG-GFP | BCH Viral Core | N/A |
| $AAV_{2/9}$-CAG-Cre-GFP | BCH Viral Core | N/A |
| **Chemicals, peptides, and recombinant proteins** | | |
| Sodium deoxycholate | Sigma-Aldrich | Cat# D6750; CAS: 160492-56-8 |
| Cycloheximide | Sigma-Aldrich | Cat# C7698; CAS: 66-81-9 |
| Heparin | Sigma-Aldrich | Cat# H3393; CAS: 9041-08-1 |
| Humulin R | Eli Lilly | Cat# U-100 |
| DTT | Sigma-Aldrich | Cat# 646563; CAS: 3483-12-3 |
| Nonidet P40 Substitute | Roche | Cat# 11 332 473001 |
| KCl | Sigma-Aldrich | Cat# P9541; CAS: 7447-40-7 |
| $MgCl_2$ | Sigma-Aldrich | Cat# 63068; CAS: 7791-18-6 |
| Tris HCl | Roche | Cat# 10812846001; CAS: 1185-53-1 |
| Percoll | Sigma-Aldrich | Cat# P1644; |

| | | |
|---|---|---|
| Corn oil | Sigma-Aldrich | Cat# C8267; CAS: 8001-30-7 |
| Tamoxifen | Sigma-Aldrich | Cat# T5648; CAS: 10540-29-1 |
| Trizol | Thermo Fisher | Cat# 15596018 |
| Dynabeads Protein A for IP | Thermo Fisher | Cat# 10002D |
| Dynabeads™ Biotin Binder | Thermo Fisher | Cat# 11047 |
| Protease inhibitor mixture | Sigma-Aldrich | Cat# P8340 |
| RNasin® Ribonuclease Inhibitors,10,000U | Promega | Cat# N2515 |
| Phosphate buffered saline | Thermo Fisher | Cat# 10010031 |
| DNase I | Sigma-Aldrich | Cat# D4527-40KU; CAS: 9003-98-9 |
| 0.05% trypsin-EDTA | Thermo Fisher | Cat# 35200-054 |
| Seahorse XF Base Medium | Agilent technologies | Cat# 102353-100 |
| Guanidine-HCl | Sigma-Aldrich | Cat# AAJ6078622; CAS: 50-01-1 |
| Beta-Amyloid (1-42), HiLyte™ Fluor 555 | Abaspec | Cat# AS-60480-01 |
| Microglia Medium (complete kit) | ScienCell Research Lab | Cat# 1901 |
| Microglia growth supplement | ScienCell Research Lab | Cat# 1952 |
| Penicillin-Streptomycin (10,000 U/mL) | Thermo Fisher | Cat# 15140-122 |
| DMEM/F-12 | Thermo Fisher | Cat# 11320-074 |
| Opti-MEM® I Reduced Serum Medium | Thermo Fisher | Cat# 31985-070 |
| Hibernate™-A Medium | Thermo Fisher | Cat# A1247501 |
| **Critical Commercial Assays** | | |
| Ultra-sensitive mouse insulin ELISA kit | Crystal Chem | Cat# 90080 |
| Mouse IGF-1 ELISA kit | RayBiotech | Cat# ELM-IGF1-1 |
| Human Amyloid beta 42 ELISA Kit | Thermo Fisher | Cat# KHB3442 |
| Murine IFN-γ TMB ELISA Development Kit | Peprotech | Cat# 900-T98 |
| TMB ELISA Buffer Kit | Peprotech | Cat# 900-T00 |
| Murine IL-4 TMB ELISA Development Kit | Peprotech | Cat# 900-T49 |
| Direct-zol RNA Microprep | Zymo Research | Cat# R2062 |
| Human Amyloid beta 1-40 ELISA | RayBiotech | Cat# ELH-AMB140-2 |
| *E. coli* Phagocytosis Assay Kit | Cayman Chemical | Cat# 601370 |
| cDNA Reverse transcription Kit | Thermo Fisher | Cat# 4368814 |
| Lipofectamine RNAiMAX Transfection | Invitrogen | Cat# 13778030 |
| Seahorse XFe24 FluxPak | Agilent | Cat# 102340-100 |
| **Experimental models: Cell lines** | | |
| Cell line: SIM-A9 microglia cells | Abm | T0247, RRID:CVCL_5I31 |
| **Experimental models: Organisms/ strains** | | |
| Moues: C57BL/6-*Tmem119*$^{em1(cre/ERT2)Gfng}$/J | The Jackson Laboratory | RRID:IMSR_JAX:031820 |
| Moues: B6J.129(Cg)-*Rpl22*$^{tm1.1Psam}$/SjJ | The Jackson Laboratory | RRID:IMSR_JAX:011029 |
| Moues: B6.Cg-Tg(APPSwFlLon,PSEN1*M146L*L286V)6799Vas/Mmjax | The Jackson Laboratory | RRID:MMRRC_034848-JAX |
| Moues: IR$^{f/f}$ | This study | Kahn lab |
| Mouse: MG-IRKO (Tmem119$^{CreERT2+/-}$::IR$^{f/f}$) | This study | N/A |
| Mouse: MG-IRKO$^{RiboTag}$ (Tmem119$^{CreERT2+/-}$::IR$^{f/f}$::RiboTag$^{HA/HA}$) | This study | N/A |
| Mouse: MG$^{RiboTag}$ (Tmem119$^{CreERT2+/-}$::RiboTag$^{HA/HA}$) | This study | N/A |
| Mouse: MG$^{IRKO}$/5xFAD (Tmem119$^{CreERT2+/-}$:: IR$^{f/f}$::5xFAD) | This study | N/A |
| **Oligonucleotides** | | |
| SMARTpool ON-TARGETplus Mouse Insr siRNA | Horizon Discovery | Cat# L-043748-00-0005 |
| SMARTpool ON-TARGETplus Mouse non-targeting control siRNA | Horizon Discovery | Cat# D-001810-01-20 |
| **Software and Algorithms** | | |
| Prism 9.0 | GraphPad | http://www.graphpad.com |
| Fiji ImageJ | NIH | https://imagej.net/software/fiji/ |
| Wave 2.6 | Agilent technologies | https://www.agilent.com/ |
| FlowJo | FlowJo | https://www.flowjo.com/ |
| ANY-maze | Stoelting | https://www.any-maze.com/ |
| R language | N/A | https://www.r-project.org/ |
| **Others** | | |
| NanoDrop 2000 | Thermo Scientific | ND2000CLAPTOP |
| Infinity Glucometer | US Diagnostics | G5-003SK |
| Magnetic Separator | Bimake | B23803 |
| Seahorse XFe24 Analyzer | Agilent technologies | https://www.agilent.com/ |
| FACS Aria™ III Sorter | BD Bioscience | https://www.bdbiosciences.com/ |
| SHIELD-based whole-brain CLARITY | LifeCanvas | https://lifecanvastech.com/ |
| RNA-sequencing | BGI Americas | https://www.bgi.com/us/home |

| | | |
|---|---|---|
| PixiMus II DEXA scan | GE Healthcare | https://www.gehealthcare.com/ |
| CLAMS (Oxymax OPTO-M3) | Columbus Instruments | https://www.colinst.com/ |

### RESOURCE AVAILABILITY

#### Lead contact

#### Materials availability

This study did not generate new unique reagents.

#### Data and code availability

- All data reported in this paper will be shared by the lead contact upon request.
- This paper does not report original code.
- Any additional information required to reanalyze the data reported in this paper is available from the lead contact upon request.

### EXPERIMENTAL MODEL

#### *Experimental animals*

All experimental procedures were conducted in accordance with the animal welfare care and were approved by the Institutional Animal Care and Use Committee (IACUC) of the Joslin Diabetes Center (JDC) and Harvard Medical School. Veterinary services are provided as needed. Mice were housed 4–5 per cage and maintained in a temperature- and humidity-controlled animal facility (23 ºC and 80% humidity) with a 12h/12h light/dark cycle (light on at 6:30 am) at the JDC. Animals were provided with standard chow diet (Diet 9F, PharmaServ) and water *ad libitum* unless otherwise specified. IR floxed mice ($IR^{f/f}$) were created and maintained at JDC. $TMEM119^{CreERT2}$ mice (Stock #031820) (Kaiser and Feng, 2019), RiboTag mice (floxed-Rpl22HA, Stock #011029) (Sanz et al., 2009) were purchased from Jackson Laboratory (Bar Harbor). MG-IRKO (**M**icro**g**lia **I**nsulin **R**eceptor **K**nock-**O**ut: $Tmem119^{CreERT2+/-}$::$IR^{f/f}$) mice were produced by crossing $IR^{f/f}$ mice and $TMEM119^{CreERT2}$ mice, to specifically delete *InsR* genes from microglia cells in a tamoxifen-inducible manner. Littermates without Cre alleles were used controls for the MG-IRKO mice. $MG\text{-}IRKO^{RiboTag}$ ($Tmem119^{CreERT2+/-}$::$IR^{f/f}$::$RiboTag^{HA/HA}$) mice were produced by crossing MG-IRKO mice with $IR^{f/f}$-$RiboTag^{HA/HA}$ mice. $MG^{RiboTag}$ ($Tmem119^{CreERT2+/-}$::$RiboTag^{HA/HA}$) mice were produced by crossing $TMEM119^{CreERT2}$ mice and RiboTag mice and served as the controls for the $MG\text{-}IRKO^{RiboTag}$ mice (age and sex matched). Mouse genotypes were verified by PCR amplification of tail extracts. Both male and female mice were included in behavioral and biochemical analyses unless otherwise specified in the figure legends.

Cre recombination was induced *in vivo* as previously reported (Garcia-Caceres et al., 2016; Sassmann et al., 2010). Tamoxifen was freshly prepared in corn oil (C8267, Sigma-Aldrich) in a final concentration of 20 mg/ml. About six-week-old mice were intraperitoneally administrated with 100 mg/kg tamoxifen (T5648, Sigma-Aldrich) once per day for consecutive 5 days. Mice were monitored daily for general health during the tamoxifen administration period. Littermates carrying *loxP*-flanked alleles but lacking expression of Cre recombinase were used as controls. Tamoxifen injected mice were recovered in cages for at least 2 weeks prior to experiments. Behavioral assays were conducted in these mice at the ages of 3- to 4-months.

#### *Isolation of primary adult mouse microglia*

Isolation of primary mouse microglia was previously described (Boroujerdi et al., 2014; Milner et al., 2022; Tamashiro et al., 2012). Briefly, we isolated primary microglia cells from 8-week male $IR^{f/f}$ mouse. Following removal of the brain and the meninges attached on the surface, brain cortices were minced in iced Hybernate A medium (A1247501, ThermoFisher). Tissues were filtered by gentle pipetting and incubated with a digestion buffer that contain a mixture of papain and DNase at 37°C for 30 min. The cell suspension was filtered through a 70 µm filter to remove myelin and cell debris. We used a 30/70 Percoll gradient centrifugation to achieve complete myelin removal. The intermediate layer of the two phases was retrieved and purified with $Cd11b^+$ beads (Miltenyi Biotech). $Cd11b^+$ cells were resuspended in microglia medium (#1901, ScienCell) in 24-well plates. The isolated primary microglia were maintained at 37°C with 5% $CO_2$/95% $O_2$. Medium was changed every other day until reaching confluence for downstream assays.

#### *Isolation of microglia using biotin-binder*

We used whole brains for microglia isolation based on a previous report (Kim et al., 2019). Briefly, indicated-age adult mice were full anesthetized and perfused with ice-cold PBS. Brain was collected and its visible vessels were removed under microscopy. Single-cell suspension was prepared followed by a gradient centrifugation (700g, 15 min, no brake) using a continuous 30% and 70% Percoll medium. Cells at the interface of the two phases was carefully collected and incubated with biotin-conjugated anti-Cd11b antibody (eBioscience, MA) on ice. The $CD11b^+$ cells were then selected using Dynabeads Biotin Binder (Invitrogen). Pellets were stored using a TRIzol reagent (Invitrogen) at -80 ºC for long-term storage, or immediately proceeded to RNA extraction (see below).

#### *Culture of SIM-A9 microglia cell line*

SIM-A9 cells, an immortalized mouse microglia cell line, was purchased from Applied Biological Materials was established from mouse cerebral cortices and have been characterized previously (Nagamoto-Combs et al., 2014). We maintained the cells in T75 flasks containing DMEM/F-12 (ThermoFisher) supplemented with 10% Fetal Bovine Serum (FBS, Sigma-Aldrich), and 100 U/mL penicillin/streptomycin (ThermoFisher) in a 37°C incubator with 5% $CO_2$/95% $O_2$. Once achieved an 80% confluence, the cells were seeded into 6-well plates through a mild incubation of 0.05% trypsin-EDTA (25300-054, ThermoFisher).

### METHOD DETAILS

#### *Blood Glucose Determination*

Blood glucose and plasma insulin levels were determined as previously described (Chen et al., 2023). Mice were subjected to 4-hr fasting, when food was withdrawn, and blood samples were collected subsequently. To measure fasting blood glucose level, a small cut on tail tip was made using a sterile surgical blade. Approximately 2 µl of venous blood was collected and blood glucose concentration was measured via a glucometer (G5-003SK, Infinity, US Diagnostics).

#### *Glucose tolerance test and insulin tolerance test*

Glucose and insulin tolerance tests were performed on 3-month-old $IR^{f/f}$ and MG-IRKO mice as described (Li et al., 2019). All tests were performed in light circle. Mice fasting for GTT and ITT procedures were allowed for free access to water at all times. For the GTT, mice were fasted overnight. The next day, individual mice were *i.p.* administered with glucose (2 mg/g/bw). For the ITT, mice fasted for 4 hrs were i.p. administered with 0.75 mIU/g body weight insulin (Humulin R; Lilly). Blood was sampled to measure glucose concentration post glucose injection.

#### *DEXA scanning and Indirect calorimetry*

DEXA scanning and CLAMS were performed as described (Almind et al., 2007; Katic et al., 2007). Briefly, 3- to 4-month-old male were anesthetized and the body composition was measured with a PixiMus II mouse densitometry system (GE Healthcare, Piscataway). For metabolic and activity analysis, mice were individually housed in indirect calorimetry chambers (Oxymax OPTO-M3 system; Columbus Instruments) and given *ad libitum* access to food and water. Within the cages, air of known $O_2$ concentration was delivered at a constant flow rate. Following a 48-hr acclimation period, the CLAMS system measured $O_2$ consumption for 60s in every 12 minutes for 24 hrs, during which food and water intake, as well as activity were measured simultaneously.

#### *Measurement of Aβ$_{42}$ and Aβ$_{40}$*

The concentration of Aβ$_{42}$ and Aβ$_{40}$ in brain homogenates was measured by Aβ$_{42}$ human ELISA kit (ThermoFisher) and Aβ$_{40}$ human ELISA kit (RayBiotech), respectively, according to manufacturer's instructions and were described previously (Chen *et al.*, 2023).

#### *Measurement of plasma insulin and IGF-1 levels*

The levels of plasma insulin and IGF-1 were measured as described previously (Chen *et al.*, 2023). Briefly, approximately 20 µL of venous blood was sampled by cutting the mouse tail tip following a 4-hr fasting. Levels of plasma insulin were measured using an ultra-sensitive mouse insulin ELISA kit (Crystal Chem), whereas levels of IGF-1 were measured using a mouse IGF-1 ELISA kit (Raybiotech), according to their respective manufacturer's instructions.

#### *siRNA transfection*

SIM-A9 cells were seeded at a density of $1 \times 10^5$ cells $cm^{-2}$ for 24 hrs prior to the transfection. The cells were transfected with 10 nM siRNA using Lipofectamine RNAiMAX (13778150, ThermoFisher) overnight in the plates according to manufacturer's protocol. The next day in the morning, culture medium was changed to fresh maintenance medium for 72 hrs until experiments. SMARTpool ON-TARGETplus control siRNA (D-001810-01-20) and mouse Insr siRNA (L-043748-00-0005) were purchased from Horizon Discovery. The target sequences for each siRNA are provided in the Supplementary Table 1. Efficiency and specificity of siRNA transfection was verified using RT-qPCR.

#### *FACS-based uptake of Aβ in primary mouse microglia*

We measured uptake of Aβ in primary mouse microglia with FACS as previously described (Chen *et al.*, 2023). IR deletion was induced in isolated microglia by infecting primary microglia with adeno-associated virus (AAV) encoding a Cre:GFP fusion protein (BCH Viral Core) for 24 hrs and cultured for an additional 5 days before experiments. Control microglia were generated using AAV encoding an empty AAV-GFP in the same experimental duration. Prior to the measurement of Aβ, IRKO microglia and the controls were changed with new culture medium and incubated with 0.1 mM Aβ$_{1\text{-}42}$ HiLyte Fluor555 (AS-60480-01, Anaspec) for 24 hrs, followed by two washes with PBS. Cells were detached from the cell culture plate and resuspended in FACS buffer (1% FBS + 0.25 mM EDTA) for FACS analysis. Aria™ III Sorter (BD) was used to gate single microglial cell and quantify the fluorescent intensity from Aβ$_{1\text{-}42}$ for individual cells.

#### *Seahorse analysis*

A Seahorse XF24 analyzer (Agilent Tech) was used to measure Oxygen Consumption Rate (OCR) in SIM-A9 microglia cells following the manufacturer's instructions based on our previous reports (Chen *et al.*, 2023; Sakaguchi et al., 2019). Briefly, cells were seeded at a density of approximately 40,000 cells per cell in an XF 24-well plate filled with 100 µL growth media. An additional 150 µL of growth media was added one hour later to a final volume of 250 µL per well. Plates were incubated in an incubator at 37 ºC overnight. We then prepared a sensor cartridge and added 1 mL of calibrant (pH 7.4) to each well and allowed for incubation at 37 ºC without $CO_2$ overnight. We removed growth media the next day and washed the cells with XF assay media. The plate was incubated at 37 ºC without $CO_2$ for 1 hr prior to the measurement. For glycolytic stress test, we loaded the following compounds respectively: glucose (10 mM), oligomycin (2 µM), 2-DG (50 mM). Results were normalized to protein content. Data were analyzed using Seahorse XF24 Wave software (v2.6).

#### *RNA Extraction and Reverse Transcription*

Total RNA was extracted from isolated microglia ($CD11b^+$) and non-microglia cells ($CD11b^{neg}$) using TRIzol (15596018, Invitrogen) according to the manufacturer's instruction. We pre-determined RNA concentration and quality using the NanoDrop 2000 (Thermo Scientific). Total RNA (0.5 µg) from each sample was reverse transcribed to cDNA using a High-Capacity cDNA Reverse transcription Kit (4368814, ThermoFisher), according to the manufacturer's user guide. We diluted the resultant cDNA libraries in ultrapure water and used diluted cDNA template (2 µL) in a 20 µl reaction system. Quantitative real time-PCR with this mixture were performed in duplicates using a SYBR Green qPCR Kit (A6002, Promega) analyzed using a CFX Connect™ System (Bio-Rad). The results were analyzed with the CFX Connect™ Software (Bio-Rad) based on the comparative CT method. All data are expressed as $2^{-\Delta\Delta CT}$ for the gene of interest normalized to the housekeeping gene *Arpp0* and presented as fold-change relative to controls. Sequences for each primer are listed in Supplementary Table 1.

#### *RiboTag isolation*

4-month-old male $MG^{RiboTag}$ and $MG\text{-}IRKO^{RiboTag}$ mice were deeply anesthetized and perfused using ice-cold PBS. The brains were separated and frozen on dry ice and stored at -80 ºC for storage. To extract microglial-specific ribosomal mRNA from bulk tissues, we performed RiboTag isolation as previously reported with modifications (Gao and Zhao, 2021; Mahadevan et al., 2020; Sanz et al., 2019). Briefly, brain hemispheres were homogenized in a pre-chilled Dounce filled in a freshly prepared iced homogenization buffer supplemented with 50 mM Tris-HCl (pH 7.5), 100 mM KCl, 12 mM $MgCl_2$, 1% NP40, 1 mM DTT, 1x protease inhibitor cocktail (B14002, Bimake), 200 U/mL RNasin (Promega), 100 µg/mL cycloheximide, and 1 mg/mL heparin. Five percent of each sample was separately prepared as *Input* samples, representing the whole transcriptome, while the remaining sample was proceeded for isolating ribosome-bound mRNA using RiboTag immunoprecipitation (*IP*) and redeemed as *IP*. mRNA from *IP* samples were subsequently isolated with a Direct-zol RNA Microprep Kit (R2062, Zymo).

#### *RNA sequencing*

Extracted mRNA samples were submitted to BGI Americas (Cambridge) as previously described (El-Darzi et al., 2022; Jiang et al., 2020). Briefly, total mRNA was first submitted to BGI Americas for quality and quantity control. After the fragmentation and reverse transcription via random primers, cDNA library was obtained. Library sequencing was performed on the DNBSEQ platform with an average generation about 4.50G Gb bases per sample. The sequencing length was PE100 (paired-end 100 bp). To obtain clean reads, the sequencing reads were filtered and stored in the FASTQ format prior to data analysis. The clean reads were aligned to mouse reference genome (Mus_musculus) by Bowtie2 on a standard reference genome version GCF_000001635.26_GRCm38.p6). Further bioinformatic analyses, including analysis of differentially expressed genes (DEG) and gene ontology (GO) biological process, were performed on BGI Americas' in-house platform *Dr. Tom* (www.bgi.com/global/dr-tom/) and a web-based analysis tool *iDep1.1* via a scandalized pipeline that was reported previously (Ge et al., 2018). We used a threshold for significantly differential expression based on experience and previous report with a fold change of ≥2 and a *P* value of ≤ 0.01.

#### *Brain homogenates preparation*

Brain homogenates were prepared according to our previous report (Chen *et al.*, 2023) with minor modifications. Briefly, following transcardially perfusion of pre-cold 1x PBS, brain tissues were cut in a sagittal manner and right hemispheres were frozen in dry ice and stored at -80ºC for biochemical analysis. To prepare lysates for ELISA measurement, we homogenized the hemispheres on ice in 5 volumes (w/v) of 0.1% Triton X-100 in TBS buffer, supplemented with protease inhibitor cocktail (B14002, Bimake) and phosphatase inhibitor cocktail (B15002, Bimake). Homogenates are ultracentrifuged at 100,000g for 1hr at 4ºC, and the resulting supernatants represented the soluble enriched fraction. The pellets than are re-suspended with 1 mL 1X RIPA extraction buffer (20-188, Millipore) supplemented with protease and phosphatase inhibitors (same as above). After sonication, homogenates were ultracentrifuged at 100,000g for 1hr at 4ºC. The resulting supernatants represented the membrane enriched fraction. The pellets were resuspended with 1mL of 5M guanidine-HCl (AAJ6078622, Fisher Scientific) solution overnight at 4ºC on a rotator and centrifuged at 20,000g for 1h at 4ºC. The resulting supernatants represented the insoluble fraction. Protein concentrations were determined using BCA protein assay kit (23227, ThermoFisher) according to instruction.

#### *Western blotting analysis*

Western blotting analysis was performed as previously described (Cai et al., 2018; Chen *et al.*, 2023). Briefly, total proteins were lysed with ice-cold RIPA lysis buffer (20-188, Millipore) containing 0.1 % SDS and a cocktail of protease and phosphatase inhibitors (Bimake, TX). Protein concentrations were quantified and normalized to the same final concentration using BCA assays. 10 to 15 ug total protein extracts were separated by electrophoresis. Following transfer, membranes were blocked for 1 hr at RT and incubated with the indicated primary antibodies overnight at 4 ºC. After three washes with 1x PBST (PBS supplemented with 0.1% Tween-20) and incubated with appropriate horseradish peroxidase (HRP)-conjugated secondary antibodies (1:10,000, Cell Signaling), membranes were visualized with SuperSignal West Pico substrate (Pierce). Chemiluminescent signals were quantified using Image J software and the band intensities were normalized to the loading control. The primary antibodies included rabbit anti-GAPDH (1:2000, #5174), rabbit anti-phospho-AMPKα1 (Thr172, 1:1000, #2535), rabbit anti-AMPKα1 (1:1000, #5831), rabbit anti-phospho-mTOR (Ser2448, 1:1000, #5536), rabbit anti-mTOR (1:1000, #2972), rabbit anti-phospho-ULK1 (Ser757, 1:1000, #14202), rabbit anti-ULK1 (1:1000, #8054), rabbit anti-beta-actin (1:1000, #4970), rabbit anti-LC3A/B (1:1000, #12741), rabbit anti-p62 (1:1000, #5114), rabbit anti-ATG3 (1:1000, #3415), rabbit anti-ATG5 (1:1000, #2630), rabbit anti-ATG7 (1:1000, #2631), rabbit anti-phospho-DRP1 (Ser637, 1:1000, #4867), rabbit anti-phospho-DRP1 (Ser616, 1:1000, #3455), rabbit anti-DRP1 (1:1000, #8570), and rabbit anti-Parkin (1:1000, #2132), all purchased from Cell Signaling Technologies. Mouse anti-MFN2 (1:1000, #ab56889), rabbit anti-citrate synthase (1:1000, #ab96600), and Total OXPHOS Rodent WB Antibody Cocktail (1:1000, ab110413) were all purchased from Abcam. Rabbit anti-FIS1 (1:1000, #sc-98900) was purchased from Santa Cruz Biotechnology.

#### *Behavioral testing*

Mouse behavioral tests were performed according to our previous reports with minor modification (Cai *et al.*, 2018; Chen *et al.*, 2023; Ferris et al., 2017). We used 4-month-old male and female mice for these behavioral tests. Each mouse was subjected to specific behavioral test only once unless otherwise stated in the legends. All mice were transferred to the behavioral testing room at least 30 min prior to the test for environmental acclimation. After each test, the behavioral equipment (arena and chamber) and associated testing items (objects and marbles) were thoroughly cleaned with 75% ethanol to eliminate potential cues left by a previous animal. Mouse behavioral performances were recorded by an HD webcam placed on the top of the behavioral arena and analyzed by ANY-maze (Stoelting).

In the open-field test (OFT), mouse was placed in an open arena (40×40×40 cm) in a position evenly lighted. During the 10-minute session, mouse was single placed in the arena for free exploration. We measured central zone entries, total distance, and average speed to assess anxiety-like behaviors.

In the tail-suspension test (TST), mouse was suspended by an adhesive tape placed 2 cm from the extremity of its tail. During the 6-minute session, immobility time was measured. In the forced-swimming test (FST), each mouse was placed in a vertical plexiglass cylinder (40 cm in height, 18 cm in diameter) containing 15 cm of water maintained at 23–25ºC. During the 6-minute session, immobility time was measured. In the sucrose preference test (SPT), the procedure was modified from a previously published protocol (Liu et al., 2018). Individual mouse was placed to a sucrose preference apparatus with ten isolated chambers (10 cm in width × 24 cm in length × 13 cm in height. There were two bottles attached on the wall of each chamber, with one filled with water and another with 1% sucrose solution (w/vol). Water and sucrose consumption were measured for a 4-hour testing session to calculate the preference score.

The marble-burying test measures obsessive compulsive-like and/or anxiety-like behaviors. We performed the procedure with minor modification as previously described (Deacon, 2006). The test was carried out in a standard testing cage (26×16×14 mm) filled to about a 5 cm-depth with wood chip. Prior to the test, twenty opaque glass marbles (14 mm in diameter) were evenly spaced on top of the bedding. Mouse was placed individually in the testing cage for a 30-min period and the number of marbles buried was recorded.

The novel object recognition (NOR) test was used to measure anxiety-like behavior. We preformed this test using a procedure modified from the previous reports (Leger et al., 2013; Lueptow, 2017). Briefly, the test was carried out in an open-field arena, similar to the OFT mentioned above. There were two phases, each consisted of a 10-min session, separated by 24 hours. During the day-1 habituation phase, mice were allowed to explore the empty arena for 10 minutes. The second phase were performed 24 hours later, which consisted of two sessions, training and novelty test, each lasted for 10 minutes. In the training session, the mouse was allowed to explore the arena with two identical objects placed in the opposite quadrants of the arena. Four hours after this session, the mouse was placed to the arena again for the novelty test. One of the objects previously visited was replaced with a novel object, consistent in height but different in shape and appearance. During the each 10-minute session, mouse behaviors were recorded and time spent exploring each object was measured to calculate discrimination index.

The light-dark (LD) box test was performed as previously descried (Salas et al., 2003) to measure anxiety-like behaviors in mice based on their innate aversion to light illuminated areas and spontaneous exploratory behavior in response to mild stressors. This test was performed in an apparatus (44×21×21 cm) containing two chambers, one bigger and bright and the other smaller and dark. During the test, a mouse was first placed in the lighted chamber and its behavioral activity was recorded for 10 minutes. The transition times between two chambers and time spent in each chamber were measured.

The social interaction (SI) test was performed as previously described with minor modification (Persico and Bourgeron, 2006). The testing arena was a three-chambered box (20×40×22 cm) made from Plexiglas. There was one rectangular opening (4×4 cm) allowing access into each chamber. The SI test consisted of 3 phases: habituation, sociability, and social novelty, each lasted for 10 minutes. The test mouse was placed in the middle chamber and allowed to explore the entire chambers. The sociability phase came after the habituation phase, during which an unfamiliar C57BL/6J male mouse (stranger 1), that had no prior contact with the test mice, was placed in one of the side chambers. The location of stranger 1 in the left vs. right side chamber was alternated between trials. The stranger mouse was enclosed in a small round wire cage (60×60×100 mm), allowing for nose-to-nose contact through the bars but prevented fighting. The animals serving as strangers had been previously habituated to placement in the small cage. During social novel phase, a second, unfamiliar mouse (stranger 2) was placed into the previously empty wire cage. Therefore, the test mouse had to make a choice between the first, already-investigated mouse (familiar stranger 1) and the novel unfamiliar mouse (new stranger 2). During each phase, the amount of time spent in each chamber and the number of entries into each chamber were recorded and analyzed by Any-Maze. SI index was calculated based on previously published protocol (Persico and Bourgeron, 2006).

#### *Whole brain clearing, staining, and imaging analysis*

Whole-brain CLARITY and imaging analysis were performed based on our previous report (Chen *et al.*, 2023) and a SHIELD protocol that was previously reported (Park et al., 2018). Post-fixed brain hemispheres were processed, imaged, and resultant 3D image datasets from light sheet microscope were analyzed, all services were provided by LifeCanvas (MA, USA). For volumetric (3D) immunofluorescent imaging, four mice from each group were processed. To prepare for brain hemispheres, fully anesthetized mice were transcardially perfused with 20 mL of ice-cold heparinized 1x PBS supplemented with 20 U/mL heparin (375095, Millipore), followed by 20 mL of ice-cold 4% PFA. After perfusion, right brain hemispheres were collected and transferred to 1x PBS supplemented with 0.02% sodium azide (4ºC) before shipping to LifeCanvas for sample clearing (SmartClear II Pro) and imaging (SmartSPIM) as described previously (Kim et al., 2015). Cleared brains were immunolabeled in SmartLabel using primary antibodies that against IBA1 (AS-60480-01, Fujifilm). A SmartSPIM light sheet microscope was used to image the brains at 3.6x with a pixel size of 1.8×1.8 µm, axial resolution of about 4 µm, and Z-step size of 4 µm. A total of 1,650 sagittal images were generated for each brain hemisphere. 3D image datasets were analyzed in SmartAnalytics to generate heat maps of IBA1 density aligned to Allen Brain Atlas (v3, 2015) (Wang et al., 2020). The results dataset contains a report on IBA1 density which was represented as cells/$mm^3$.

#### *Immunohistochemistry staining*

Brain tissues were fixed in 4% PFA overnight at 4ºC. 24 hrs later, tissues were dehydrated in 30% sucrose in 1x PBS for another 48 hrs and cut at a 40-µm thickness using a cryostat. Primary antibodies that were used for immunohistochemistry staining were: Iba1 (1:1000, #019-19741, Wako), β-amyloid (1:500, ab201060, Abcam), goat anti-rabbit IgG conjugated to Alexa Fluor 488 (1:500, ab11008, Abcam). Sections were incubated with selected primary antibodies overnight at 4ºC with gentle shaking, followed by incubation with the secondary antibody at RT for 1 hr and coverslipped with DAPI for counterstaining. Slides were scanned and analyzed using Image J.

#### *Transmission electron microscopy (TEM)*

TEM was performed as previously described with modification (Kim *et al.*, 2019; Viana-Huete et al., 2018). Briefly, 3-month-old $IR^{f/f}$ and MG-IRKO male mice were anesthetized and transcardially perfused with fixative containing 4% PFA (Electron Microscopy Tech) and 2.5% glutaraldehyde grade I in 0.1 M sodium phosphate buffer (pH 7.3) overnight at 4°C. Tissue blocks of approximately 8 $mm^3$ were collected from selective brain areas and subjected to Joslin Advanced Microscopy Core for EPON embedding by standard protocols. Samples were postfixed in 1% OsO4, 1.5% $K_4[Fe(CN)_6]$, dehydrated with acetone, and embedded in epon-812 (Taab). Ultrathin sections (60 nm) in the hypothalamus were obtained with an Ultracut E ultramicrotome (Leica), stained with lead citrate, and examined under a JEOL 1011 TEM with a Hamamatsu Orca9 HR Digital Camera at the Core in a blind manner to the groups. The features of microglia were identified as previously reported (Garcia-Cabezas et al., 2016). Images were generated from TEM at a magnification of ×15,000. For quantification, we measured the number and mitochondria area from at least 10 randomly selected microglial cells per animal.

#### *Quantification and Statistical Analysis*

All statistical analysis was performed using Prism 9 GraphPad (San DiegoSample size was estimated based on experience from previous work from our and other groups. Two-way ANOVA analysis was used to determine the effects of two distinct variables, such as genotype and treatment. For repeated measure analysis, ANOVA was used when values were over different times. Post-hoc analyses were used when appropriate and are depicted in each figure legend. For comparison between only two groups, statistical significance was determined by an unpaired Student's *t*-test. A value of $P < 0.05$ was considered statistically significant. All data are shown as mean ± SEM unless otherwise stated.

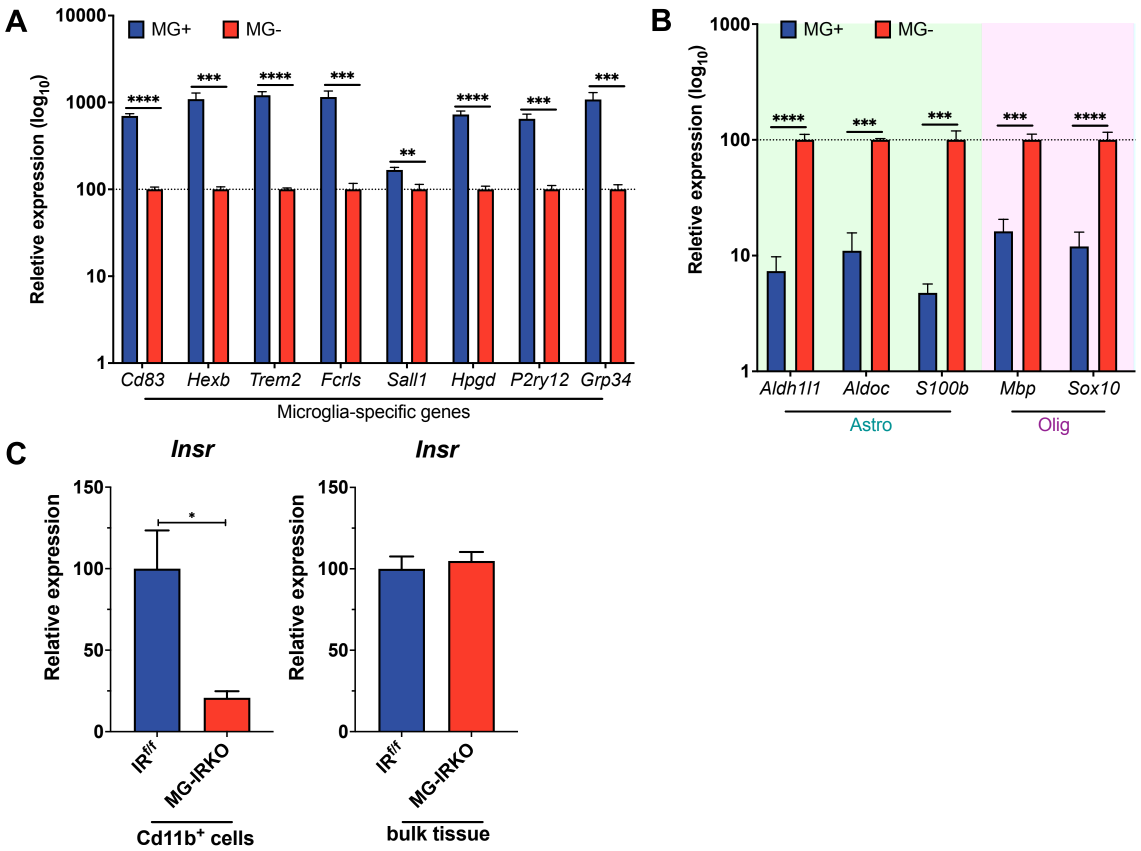


**Figure S1**. ***Deletion of InsR in Microglia, Related to Figure 1***

(A-B) RT-qPCR analysis showing relative mRNA levels (on a $\log_{10}$ scale) of signature genes of (A) microglia and (B) astrocyte and oligodendrocytes in isolated microglia or non-microglia cells. N = 6 mice per group. (C) RT-qPCR analysis of relative mRNA levels of *Insr* in isolated microglia from whole-brain (left) and in bulk cortex tissues (right) from male $IR^{f/f}$ control and MG-IRKO mice (3-month-old, male). N = 4-6 per group. *Arpp0* was used as the housekeeping control. Data are mean ± SEM. *$P < 0.05$, **$P < 0.01$, ***$P < 0.001$, ****$P < 0.0001$, unpaired *t*-test.

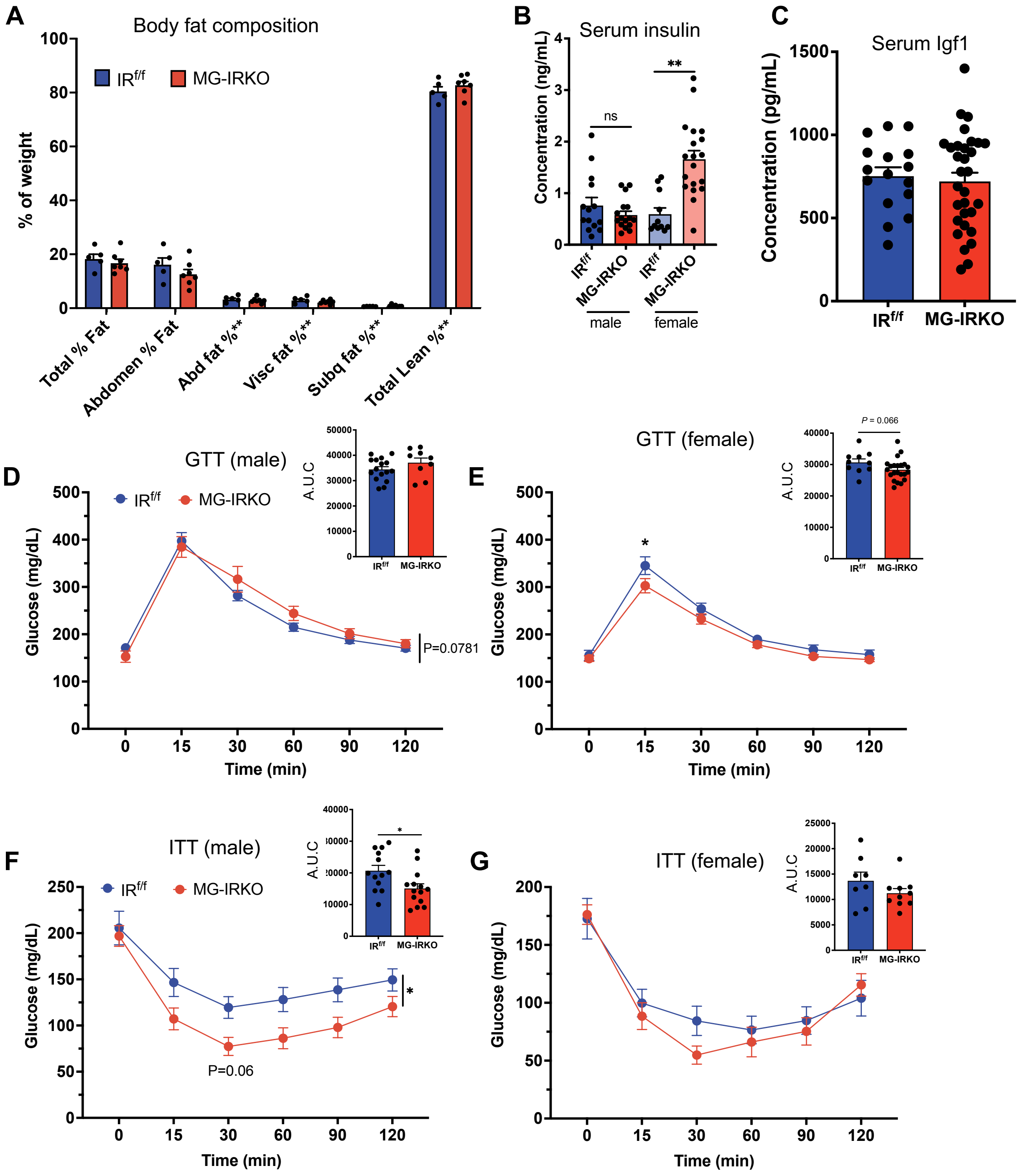


**Figure S2**. ***Deletion of InsR in Microglia had minimal effects on body fat composition and glucose/insulin sensitivity, Related to Figure 1***

(A) body fat composition measured by DEXA scanning in male IR$^{f/f}$ control and MG-IRKO mice (N = 5-7 per group). (B) serum concentration of insulin in IR$^{f/f}$ and MG-IRKO mice on 4-hr fasting (N = 16-31 per group). (C) serum concentration of Igf1 in $IR^{f/f}$ and MG-IRKO mice on 4-hr fasting (N = 16-31 per group). $^{**}P < 0.01$, unpaired *t*-test. (D-E) glucose tolerance test in (D) male and (E) female mice, N = 9-22 per group. (F-G) insulin tolerance test in (D) male and (E) female mice, N = 8-15 per group. $^{*}P < 0.05$, two-way RM ANOVA followed by Sidak's multiple comparisons test. Data are mean ± SEM.

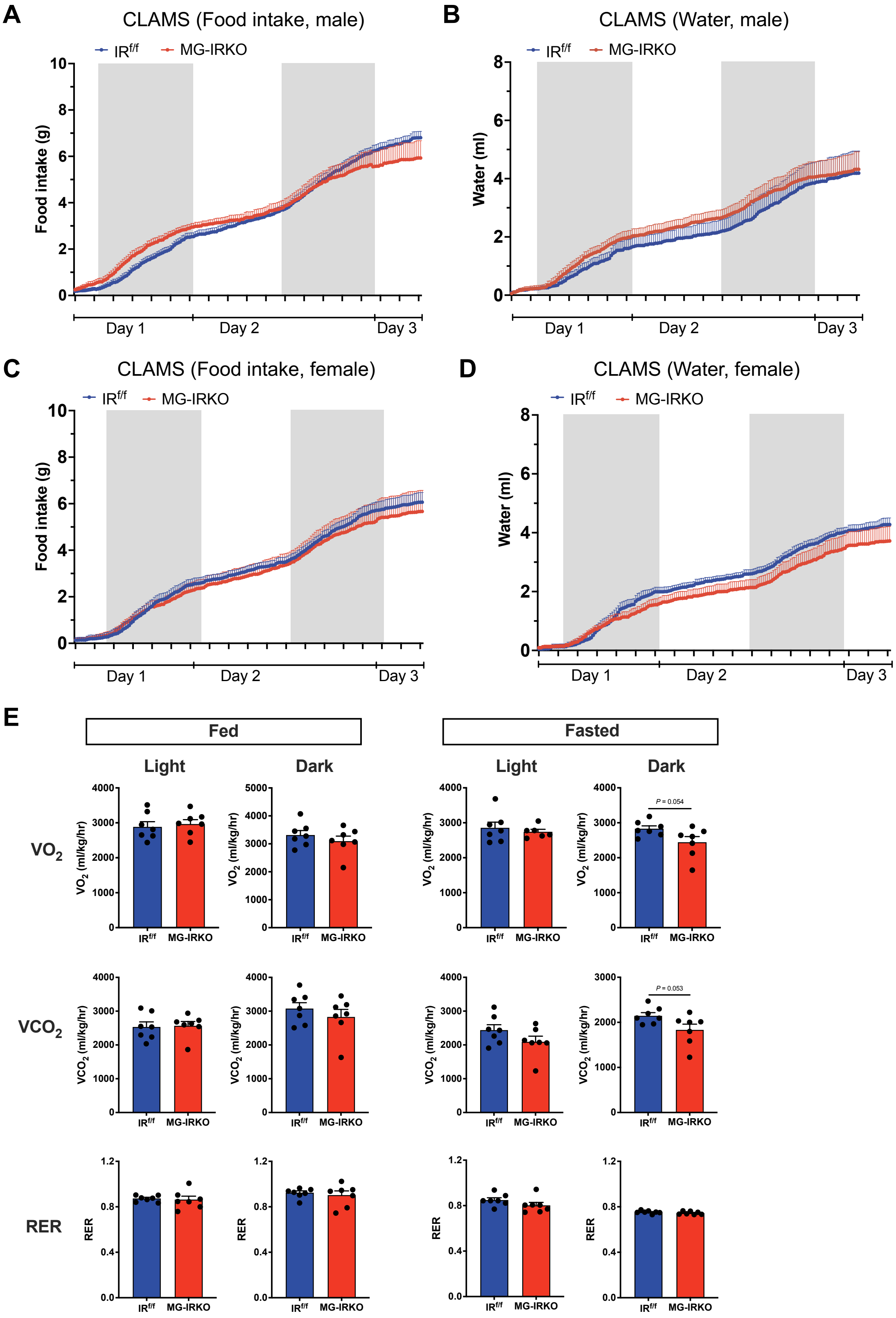
A
CLAMS (Food intake, male)
B
CLAMS (Water, male)
C
CLAMS (Food intake, female)
D
CLAMS (Water, female)
IR$^{f/f}$
MG-IRKO
Food intake (g)
Water (ml)
Day 1
Day 2
Day 3
E
Fed
Fasted
Light
Dark
VO$_2$
VCO$_2$
RER
VO$_2$ (ml/kg/hr)
VCO$_2$ (ml/kg/hr)
P = 0.054
P = 0.053

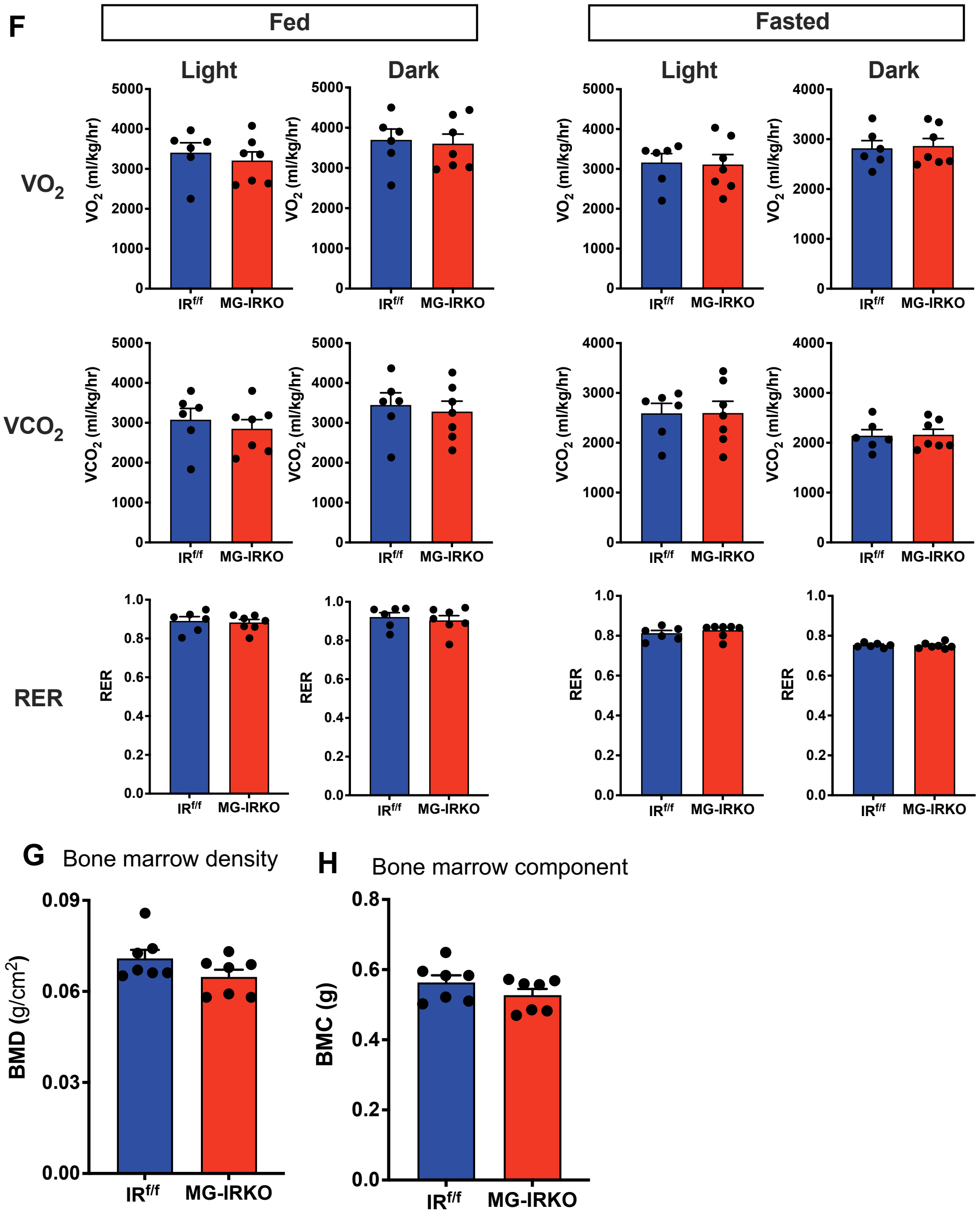


**Figure S3**. ***Metabolic phenotype in mice with microglia IR deletion, Related to Figure 1***

(A-D) food and water intake in 4-month-old (A-B) on-fed male and (C-D) female mice (N = 7 per group). Shared area indicates dark period during the testing day. (E-F) $O_2$ and $CO_2$ composition and RER measured by CLAMS in 4-month-old (E) male and (F) female mice (N = 7 per group) on both fed and fasted states. (G) bone marrow density and (H) bone marrow component in 4-month-old male mice (N = 7 per condition). Data are mean ± SEM. Two-way RM ANOVA.

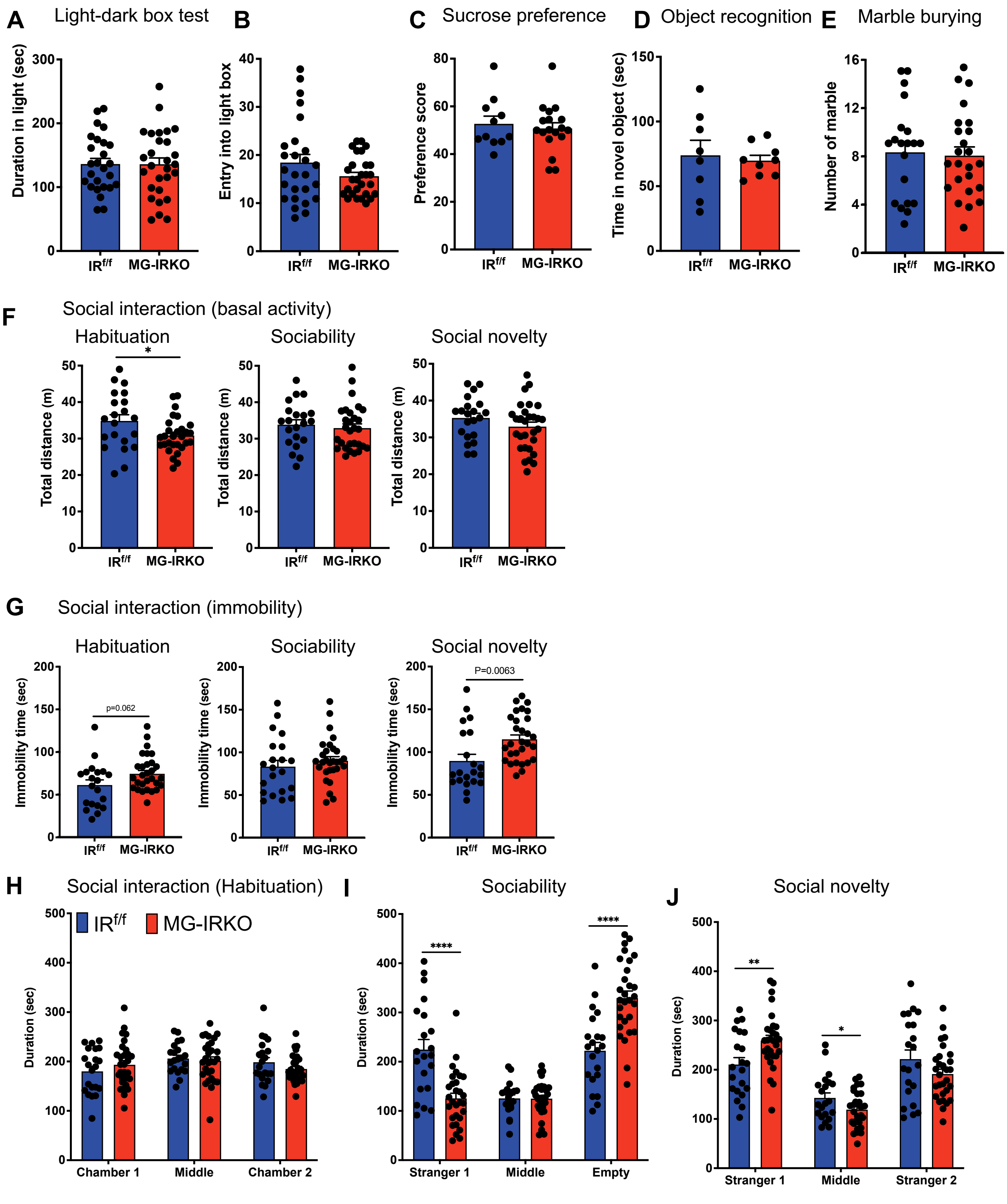


**Figure S4**. ***Effects of Microglia IRKO on behavioral performances, Related to Figure 1***

(A-B) Performance of 3-month $IR^{f/f}$ and MG-IRKO male mice in the light-dark box test, showing (A) duration in light (sec) and (B) entry into light box. N = 26-28. (C) Preference score of 3-month $IR^{f/f}$ and MG-IRKO male mice in the sucrose preference test. N = 11-18; (D) Time spent on novel object of 3-month $IR^{f/f}$ and MG-IRKO male mice in the novel object recognition test. N = 8-9; (E) Number of marbles buried of 3-month $IR^{f/f}$ and MG-IRKO male mice in the marble burying test. N = 8-9; (F-H) Performance of 3-month $IR^{f/f}$ and MG-IRKO male mice during the three sessions in the social interaction test, showing (F) total distance (m), (G) immobility time (sec) and (H) duration of time spent in individual chambers. N = 22-29. Data are mean ± SEM. *$P < 0.05$, **$P < 0.01$, ****$P < 0.0001$, unpaired *t*-test.

**A** OXPHOS pathway genes

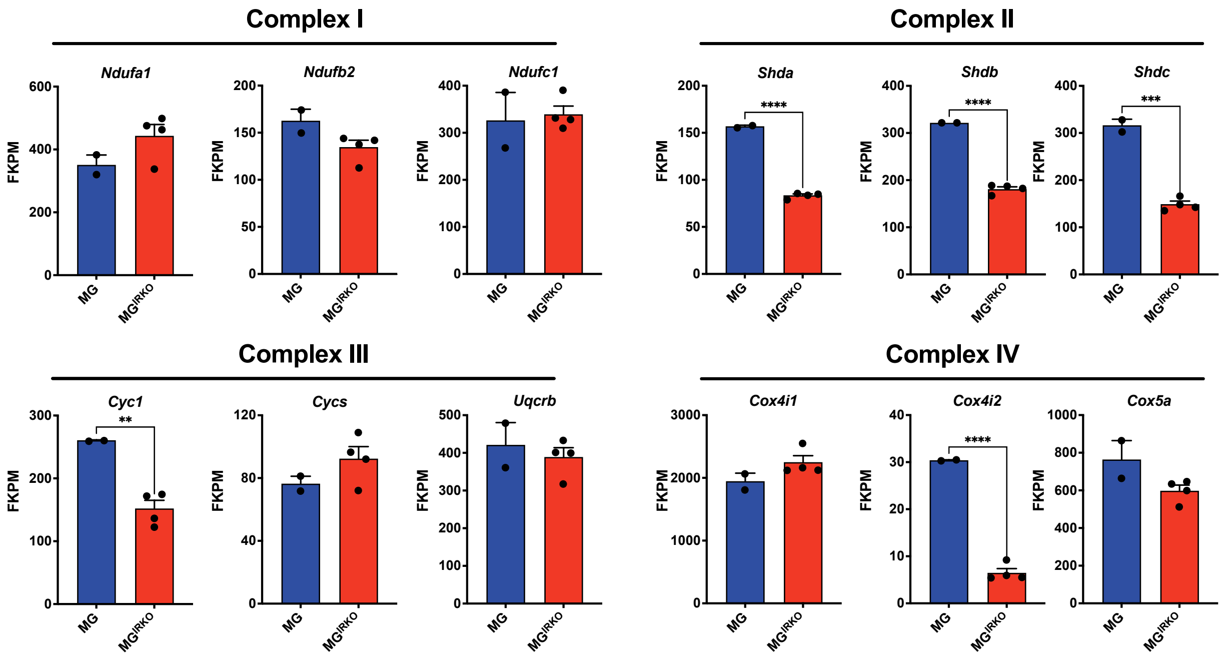


**Figure S5**. ***RiboTag reveals that microglia IR deletion impairs OXPHOS pathway, Related to Figure 2***

(A) FKPM values of example list of genes encoding pathways of mitochondrial complex I-V. N = 2-4. Data are mean ± SEM. ***P* < 0.01, ****P* < 0.001, *****P* < 0.0001, unpaired *t*-test.

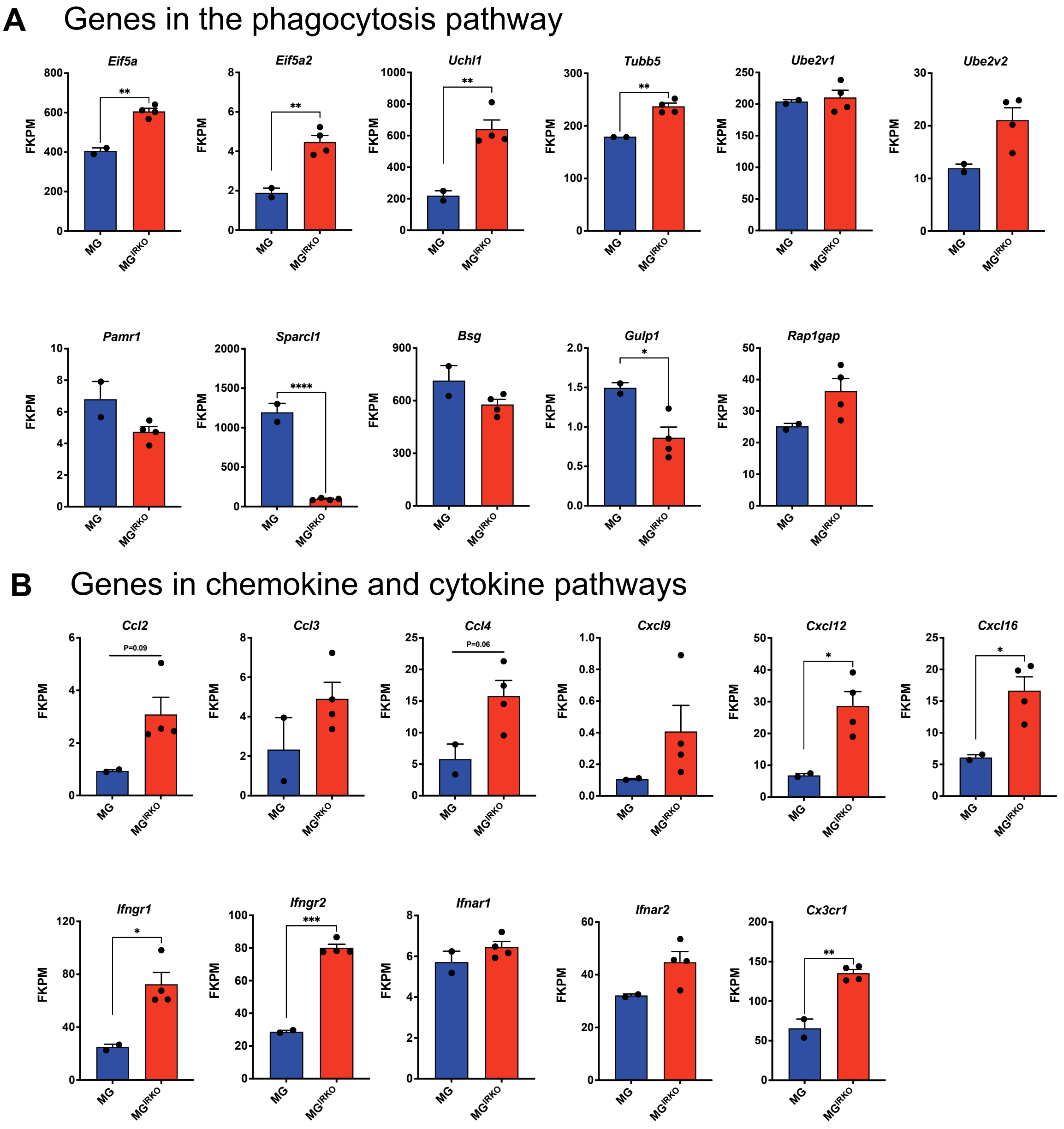


**<u>Figure S6</u>. *RiboTag reveals that microglia IR deletion activates immune pathways, Related to Figure 3***

(A-B) FKPM values of example list of genes from RiboTag profiling, including genes encoding pathways of (A) phagocytosis and (B) chemokine and cytokine. N = 2-4. Data are mean ± SEM. *$P$ < 0.05, **$P$ < 0.01, ***$P$ < 0.001, unpaired $t$-test.

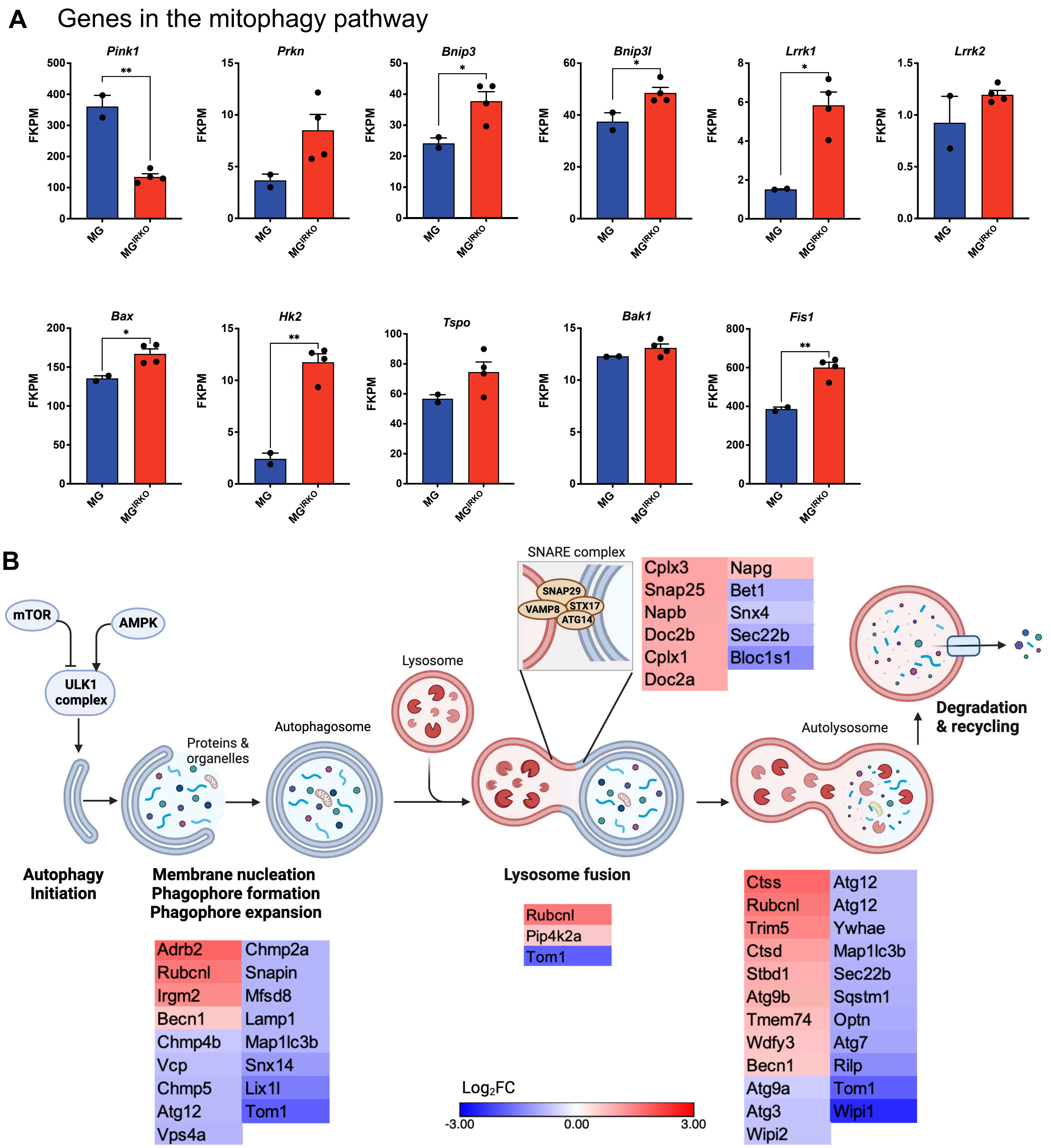


**Figure S7. *Microglia IR deletion impairs autophagy and mitophagy, Related to Figure 3***

(A) FKPM values of example list of genes of mitophagy from RiboTag profiling. N = 2-4. Data are mean ± SEM. **P* < 0.05, ***P* < 0.01, unpaired *t*-test. (B) Schematic and heatmap showing DEGs involved in the regulation of autophagy. The magitude of regulation is represented in the heatmap by $log_2$FC value (MR-IRKO / $IR^{f/f}$ control).

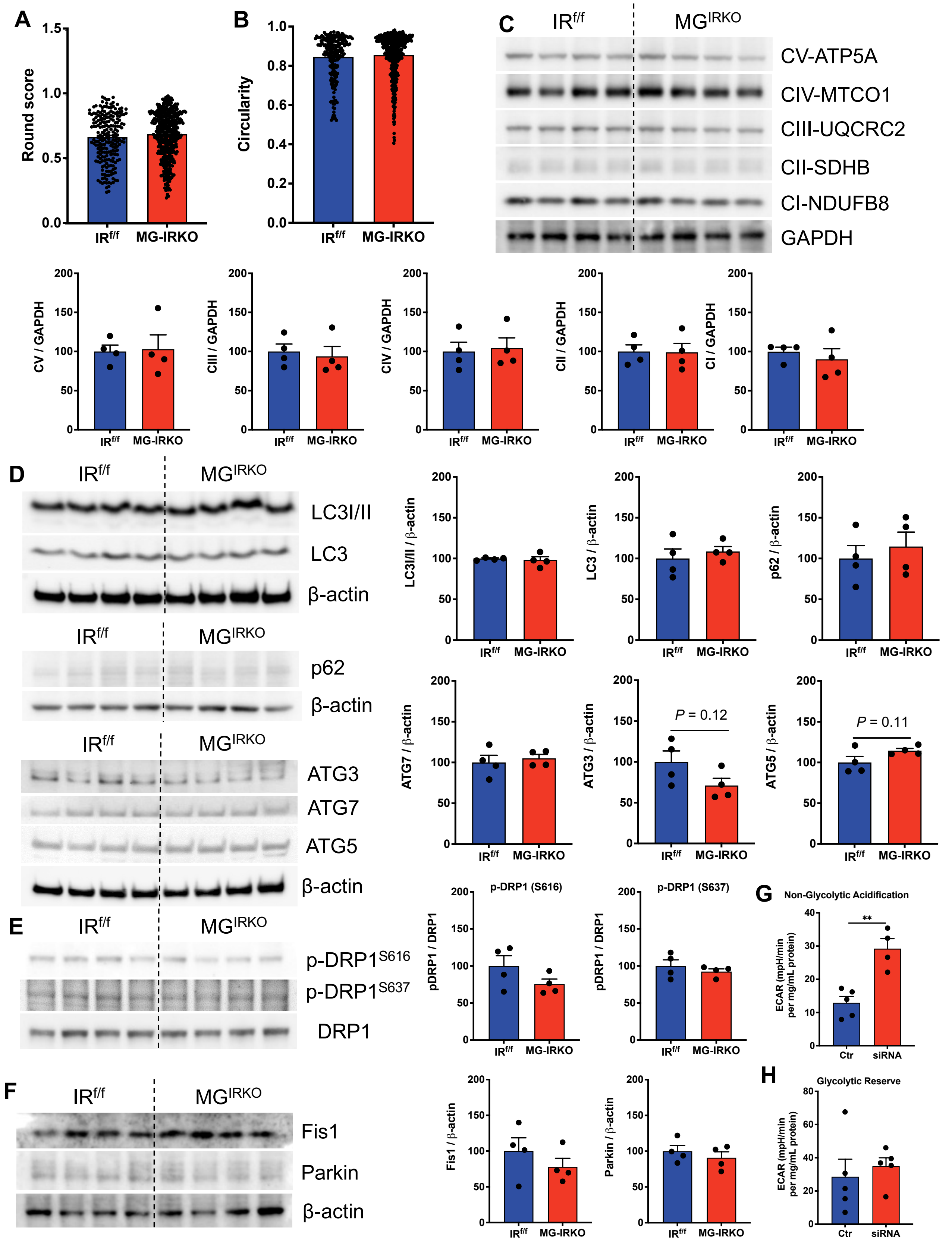

A
Round score
IRf/f
MG-IRKO
B
Circularity
C
IRf/f
MGIRKO
CV-ATP5A
CIV-MTCO1
CIII-UQCRC2
CII-SDHB
CI-NDUFB8
GAPDH
CV / GAPDH
CIII / GAPDH
CIV / GAPDH
CII / GAPDH
CI / GAPDH
D
LC3I/II
LC3
β-actin
p62
ATG3
ATG7
ATG5
LC3I/II / β-actin
LC3 / β-actin
p62 / β-actin
ATG7 / β-actin
ATG3 / β-actin
ATG5 / β-actin
P = 0.12
P = 0.11
E
p-DRP1S616
p-DRP1S637
DRP1
p-DRP1 (S616)
p-DRP1 (S637)
pDRP1 / DRP1
F
Fis1
Parkin
Fis1 / β-actin
Parkin / β-actin
G
Non-Glycolytic Acidification
ECAR (mpH/min per mg/mL protein)
Ctr
siRNA
H
Glycolytic Reserve

**Figure S8**. ***Loss of microglia IR signaling impairs cellular metabolism, Related to Figure 4***

(A-B) TEM analysis of mitochondria morphology. Bar graph showing mitochondrial roundness score (A) and circularity index (B). N = 48 microglia from N = 4 $IR^{f/f}$ male mice, and N = 57 microglia from N = 4 MG-IRKO male mice. (C-F) Immunoblot analysis of (C) mitochondrial complex proteins, (D) autophagy protein, (E) phosphorated DRP1 proteins and their total protein DRP1, and (F) mitophagy proteins in brain hippocampal homoganates from $IR^{f/f}$ and MG-IRKO mice (4-month-old, male). N = 4. $^{**}P < 0.01$, unpaired *t*-test. The protein levels were normalized to GAPDH or beta-actin or respective total proteins. Note that the reference protein GAPDH band was shared in Figure 4D. (G-H) Seahorse analysis of IRKD microglia subjected to glycolysis stress, showing non-glycolytic acifification (G) and glycolytic reserve (H). N = 4-5. $^{**}P < 0.01$, two-way RM ANOVA analysis. Data are mean ± SEM.

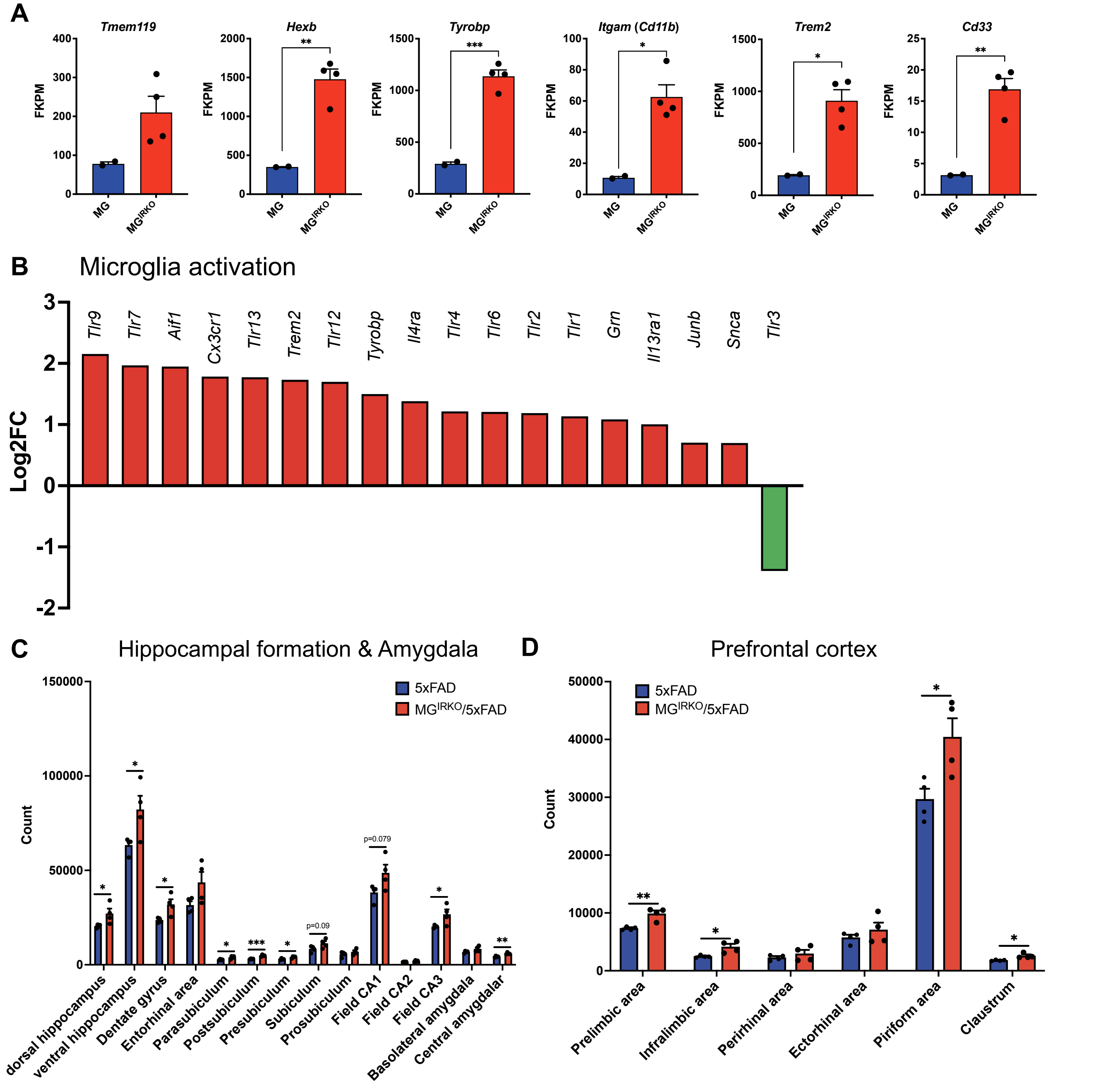


***Figure S9. Loss of microglia IR signaling causes microglia activation and impairs Aβ uptake. Related to Figure 5***

(A) FKPM value of selected microglia gene markers, including *Tmem119*, *Hexb*, *Tyrobp, Itgam*, *Trem2*, *Cd33*. (B) DEGs involved in the regulation of microglia activation are represented by $Log_2FC$. (C-D) SHIELD-based CLARITY analysis of IBA1 expression in (D) hippocampal formation and amygdala and

(E) prefrontal cortex from $MG^{IRKO}$/5xFAD mice compared to 5xFAD mice. N = 4. Data are mean ± SEM. *$P$ < 0.05, **$P$ < 0.01, ***$P$ < 0.001, unpaired $t$-test.

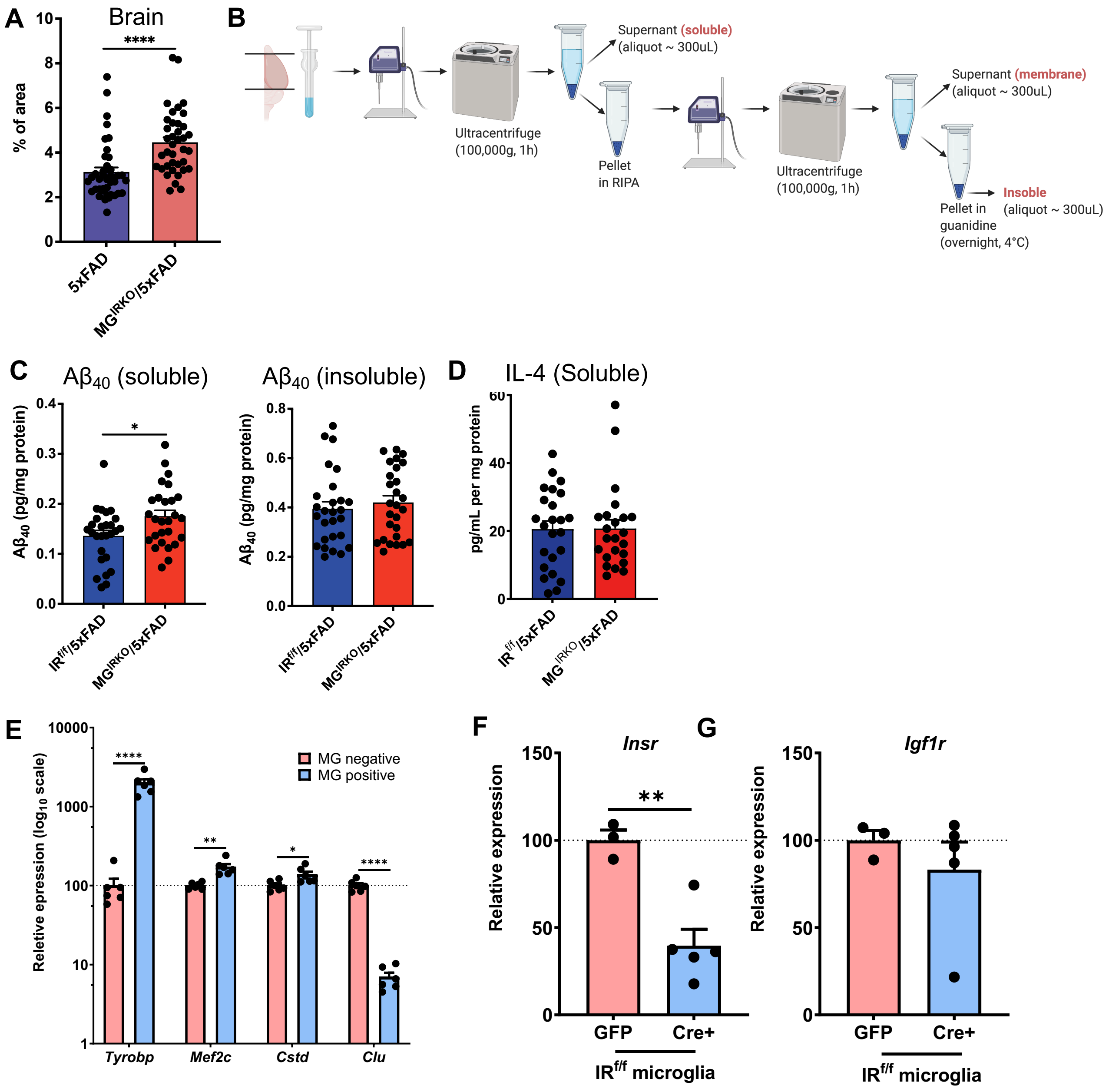


### Figure S10. *Loss of microglia IR signaling exaggerates AD pathology. Related to Figure 6*

(A) IHC analysis of β-amyloid protein in the sagittal brain sections cut from 6-month male MG$^{IRKO}$/5xFAD mice compared to 5xFAD mice, bar graph showing % of area in the brain sections. N = 4 mice per group. (B) Procedures of extraction of soluble and insolube fractions of brain hemespheres. (C) ELISA for Aβ$_{40}$ levels in soluble and insoluble fractions from brain homogenates of 6-month 5xFAD and MG$^{IRKO}$/5xFAD mice; N = 27 mice per group. (D) ELISA for IL-4 levels in soluble fraction from brain homogenates of 6-month 5xFAD and MG$^{IRKO}$/5xFAD mice; N = 27 mice per group. (E) qPCR analysis of selected microglia signature genes in relative expression ($log_{10}$ scale) in microglia positive ($Cd11b^+$ cells) and microglia negative cells ($CD11b^-$ cells) isolated from male adult $IR^{f/f}$ mice. N = 6. (F-G) Relative expression of mRNA of (F) *Insr* and (G) *Igf1r* in purified microglia isolated from male adult $IR^{f/f}$ mice and infected with AAV-Cre:GFP. N = 3 – 5. Data are mean ± SEM. **P* < 0.05, ***P* < 0.01, *****P* < 0.0001, unpaired *t*-test.
